# Engineered Flt3L Drives Tolerogenic State to Attenuate Anti-drug Antibody Responses

**DOI:** 10.1101/2024.03.21.586168

**Authors:** Aaron T. Alpar, Rachel P. Wallace, Kirsten C. Refvik, Suzana Gomes, Ani Solanki, Laura T. Gray, Anna J. Slezak, Abigail L. Lauterbach, Lauren A. Hesser, Shijie Cao, J. Emiliano Gómez Medellín, Lauren G. Robinson, Jeffrey A. Hubbell

**Affiliations:** Pritzker School of Molecular Engineering, University of Chicago; Chicago, IL 60637, USA; Animal Resources Center, University of Chicago; Chicago, IL 60637, USA; Department of Pharmaceutics, University of Washington; Seattle, WA 98195, USA.; Department of Biomedical Engineering, Johns Hopkins University; Baltimore, MA 21218, USA; Committee on Immunology, University of Chicago; Chicago, IL 60637, USA; Committee on Cancer Biology, University of Chicago; Chicago, IL 60637, USA

## Abstract

Immune reactions to protein drugs present substantial challenges to protein replacement for treating congenital diseases and metabolic deficiencies, due to the lack of endogenous tolerance or the protein drug’s partial or total non-human origin. We sought to transiently modify the immune environment when the adaptive response to the drug antigen is mounted to lessen future reactions upon continued therapeutic treatment, without modifying the drug itself. Herein, we characterize a recombinant fusion of the cytokine Flt3L to serum albumin and describe a novel pathway of Flt3L-mediated immune regulation. We highlight reduced activation of dendritic cells (DC) as well as an increased frequency of DCs expressing LAP, a TGF-β precursor. These effects in combination with low doses of the exogenous antigen led to less TH2 differentiation. This enabled a tolerance-biasing induction regimen to significantly decrease anti-drug antibodies upon repeated exposure to a clinically used, immunogenic fungal enzyme, rasburicase. This induction regimen reduced the Tfh compartment and increased Tfh cells expressing Foxp3 and PD-L1, suggesting a regulatory response. Overall, we introduce the use of a Flt3L variant as an induction therapeutic to modulate the innate immune response, thereby attenuating the adaptive reaction to antigenic protein drugs and addressing an unmet clinical need.

## Introduction

Biologics are the preferred treatment for several health conditions, including cases of enzyme replacement.(1, 2) Because these drugs are commonly foreign proteins, novel fusions, or replace a genetically-absent protein, the patient can react and mount an immune response to the drug, leading to a high rate of treatment failure.(3–6) Such failure is usually due to either inhibition of the drug function or the development of infusion reactions, both of which are mediated by a patient’s antibody response to the biologic.(7–11) Current clinical methods to ameliorate the development of anti-drug antibodies (ADAs) include administering immunosuppressive drugs such as methotrexate or rituximab; however, these options leave the patient susceptible to secondary infection. Previous work in the Hubbell lab has sought to modify the initial presentation of the antigen to promote tolerogenic mechanisms.(12–16) Additionally, other groups have investigated administration of low dose antigen in a tolerogenic regimen with other immunosuppressing drugs,(17) such as encapsulated rapamycin,(18–20) or in nanoparticles comprising phosphatidyl serine to induce tolerogenic APC uptake,(21) the latter of which has proven beneficial in reduction of ADA formation and specifically the formation of “inhibitors” in the case of hemophiliacs. Indeed, previous work has demonstrated that a treatment regimen using a dendritic cell (DC) growth factor, FMS-like Tyrosine kinase 3 Ligand (Flt3L), in combination with rapamycin and low dose antigen can attenuate ADAs against Factor VIII, demonstrating the importance of the antigen presenting compartment in initiating and perpetuating the anti-drug immune response.(19)

DCs have become increasingly relevant in tolerance induction strategies as their functions beyond antigen presentation are elucidated. Acting as regulators between pro- and anti-inflammatory cascades, DCs can dictate the trajectory of immune reactions as the production of cytokines and upregulation of surface receptors enables crosstalk with the adaptive immune response.(22–29) As to pro-inflammatory responses, type 1 conventional DCs (cDC1) play an important role in the anti-tumor response. By cross-presenting exogenous antigens to cytotoxic CD8^+^ T cells, cDC1s educate these T cells and stimulate inflammatory immune reactions against tumors and infections.(30–36) Enhancing cDC1 presence in the tumor microenvironment via chemokine localization or through expansion of these cells has shown therapeutic benefit in combination with other immunostimulatory agents, such as checkpoint blockades, irradiation, or addition of therapeutic adjuvants.(37–40) Expansion of DCs has been achieved through the use of irradiated tumor cells acting as factories of GM-CSF (GVax) or Flt3L (FVax) or through exogenous addition via regular injections of the recombinant protein.(37, 38, 41–47)

DCs can also act in immunoinhibitory capacities.(48, 49) The ability of DCs to restrain an immune reaction is understudied in contrast to research focusing on the stimulatory ability of these cells. One such model is the research into mregDCs, a subset of mature DCs enriched in immunoregulatory molecules.(50–53) While these cells maintain antigen presentation capacity, their co-expression of regulatory markers such as CD200 and PD-L1 inhibits inflammatory activation of adaptive cells. Other groups have described tolerogenic mechanisms by which DCs are able to promote tolerance and Treg formation through the production of IL-10, indoleamine 2,3-dioxygenase (IDO), TGF-β, and retinoic acid, as well as protocols to generate such tolerogenic DCs as a therapeutic.(54–60)

In this work—in contrast to previous studies focusing on antigen modification—we engineered a recombinant fusion of the DC growth factor Flt3L to serum albumin (SA) to modulate the antigen-presenting compartment and bias it toward tolerance. Other research has demonstrated the utility of Flt3L in preventing immune reactions; however, this was performed in combination with other immunosuppressive agents such as rapamycin and required repeated Flt3L infusions.(19) To this end, fusion to SA improves the overall pharmacologic properties of Flt3L and reduces the burden of frequent administration. Additionally, we describe the cellular pharmacodynamic response to Flt3L-SA, which aligns with other work using Flt3L-albumin fusions.(40, 61) We examined the phenotype of the generated DCs and probed the expansion of regulatory T cells (Tregs), a known effect of Flt3L treatment with contentious ontogeny(48, 62–64). Additionally, we describe decreased activation of the generated DCs, hallmarked by reduced expression of MHCII and PD-L1, as well as expansion of a TGF-β-expressing DC population, accompanied by an increase in Treg numbers. When two low doses of exogenous antigen were co-delivered with Flt3L-SA, Treg and TH2 differentiation was reduced. Additionally, in a murine model using enzyme replacement therapy, we observed a reduction in ADA development and anaphylaxis upon repeated high-dose exposure to the foreign biologic, accompanied by a reduction in antigen-specific germinal center (GC) B cells and an increase in T follicular regulatory (Tfr) cells, including those expressing PD-L1.

## Results

### Serum albumin fusion minimally impacts Flt3L bioactivity

To enhance the circulation properties of murine Flt3L, we recombinantly fused murine serum albumin (SA) to the C terminus of murine Flt3L, separated by a (G_4_S)_2_ linker (Figure 1A).(65, 66) Both the native, soluble Flt3L (WT) and the SA fusion (Flt3L-SA) were produced in-house prior to affinity purification and size exclusion chromatography. After purification, pure Flt3L variants were observed, including glycoforms in the WT protein and a minor dimeric population in Flt3L-SA (Figure 1B). Additionally, both WT and Flt3L-SA show similar affinity (K_D_=160 pM) for their cognate receptor CD135, as measured by ELISA (Figure 1C). To test the bioactivity of the variants, we stimulated bone marrow-derived dendritic cells (BMDCs) and measured ERK1/2 phosphorylation. Via western blot, peak ERK1/2 phosphorylation was observed following 5 min of cytokine exposure (Figure 1D). By flow cytometry, we then determined that the EC_50_ of the WT Flt3L was 1.3 nM, while that of Flt3L-SA was 3.6 nM (Figure 1E). We take these data to demonstrate that both recombinant WT Flt3L and Flt3L-SA are bioactive.

**Figure 1:**
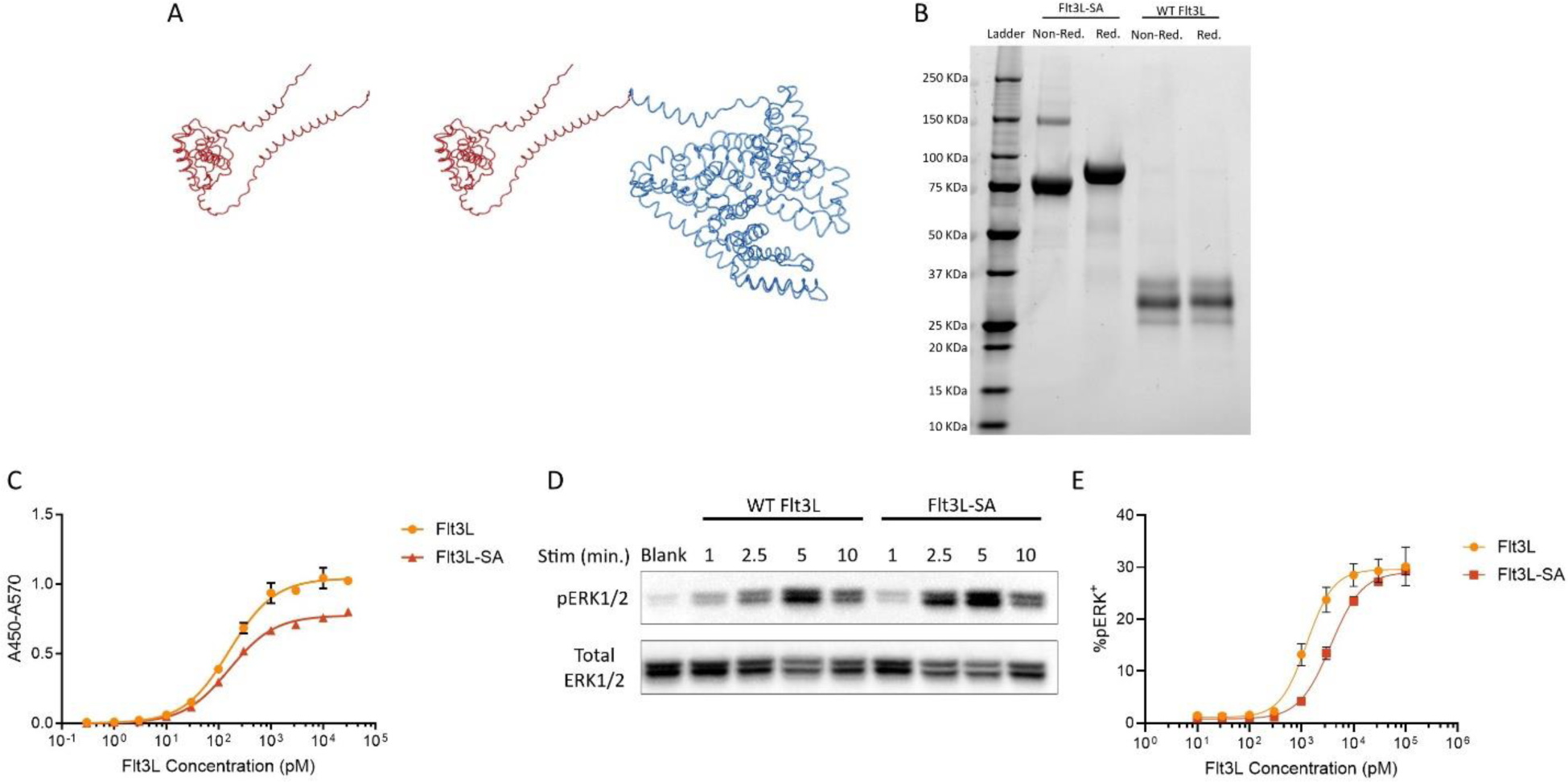
Biochemical characterization of Flt3L variants. (a) Representation of the WT (left) and the C-terminal SA fusion (right, Flt3L in red and serum albumin in blue) variants of Flt3L generated using AlphaFold and formatted in BioRender. (b) SDS-Page separation of purified Flt3L-SA under non-reducing (lane 2) and reducing (lane 3) conditions as well as the WT Flt3L under non-reducing (lane 4) and reducing (lane 5) conditions. (c) Affinity for each of the constructs for the receptor CD135 (Flt3) via ELISA. Affinities were measured as 160.4±10.7 pM and 160.0±7.3 pM for WT Flt3L and Flt3L-SA, respectively, via non-linear fit for one site specific binding reported as K_D_ ± standard deviation. (d) Bioactivity of each variant as assessed over time via ERK1/2 phosphorylation on serum-starved, Flt3L-generated BMDCs. Cells were stimulated for the indicated number of minutes at a concentration of 2 µM of WT Flt3L or Flt3L-SA before lysis and evaluation by Western blot. (e) Flt3L shows an EC50 of 1.3 nM, while Flt3L-SA has an EC50 of 3.6 nM as assessed by phosphorylation of ERK1/2 via flow cytometry after five-minute stimulation on cells generated as in figure 1D using a sigmoidal non-linear fit.

### Serum albumin fusion enhances the pharmacokinetic properties of Flt3L

We next sought to determine how the fusion to serum albumin influences the half-life and circulation of Flt3L-SA in comparison to the WT protein. Mice were subcutaneously (s.c.) injected with 10 µg of either of the two variants, based on Flt3L equivalent mass, and blood was collected at the timepoints indicated (Figure 2A). A marked increase in plasma Flt3L was observed in mice treated with Flt3L-SA at all measured timepoints, demonstrating prolonged kinetics and increased availability of the cytokine (Figure 2B). Integration of this curve with respect to time revealed that mice treated with Flt3L-SA had a roughly 30-fold increase in total Flt3L exposure as compared to those treated with the WT form (Figure 2C). Furthermore, the half-life of WT Flt3L was extended more than fivefold by the Flt3L-SA fusion (10.6 hr vs. 55.8 hr), and the time to reach maximal levels was delayed from 1.3 hr to 48 hr (Figure 2D). These pharmacokinetic values are comparable to those seen for other Flt3L-serum albumin fusion proteins.(40)

**Figure 2:**
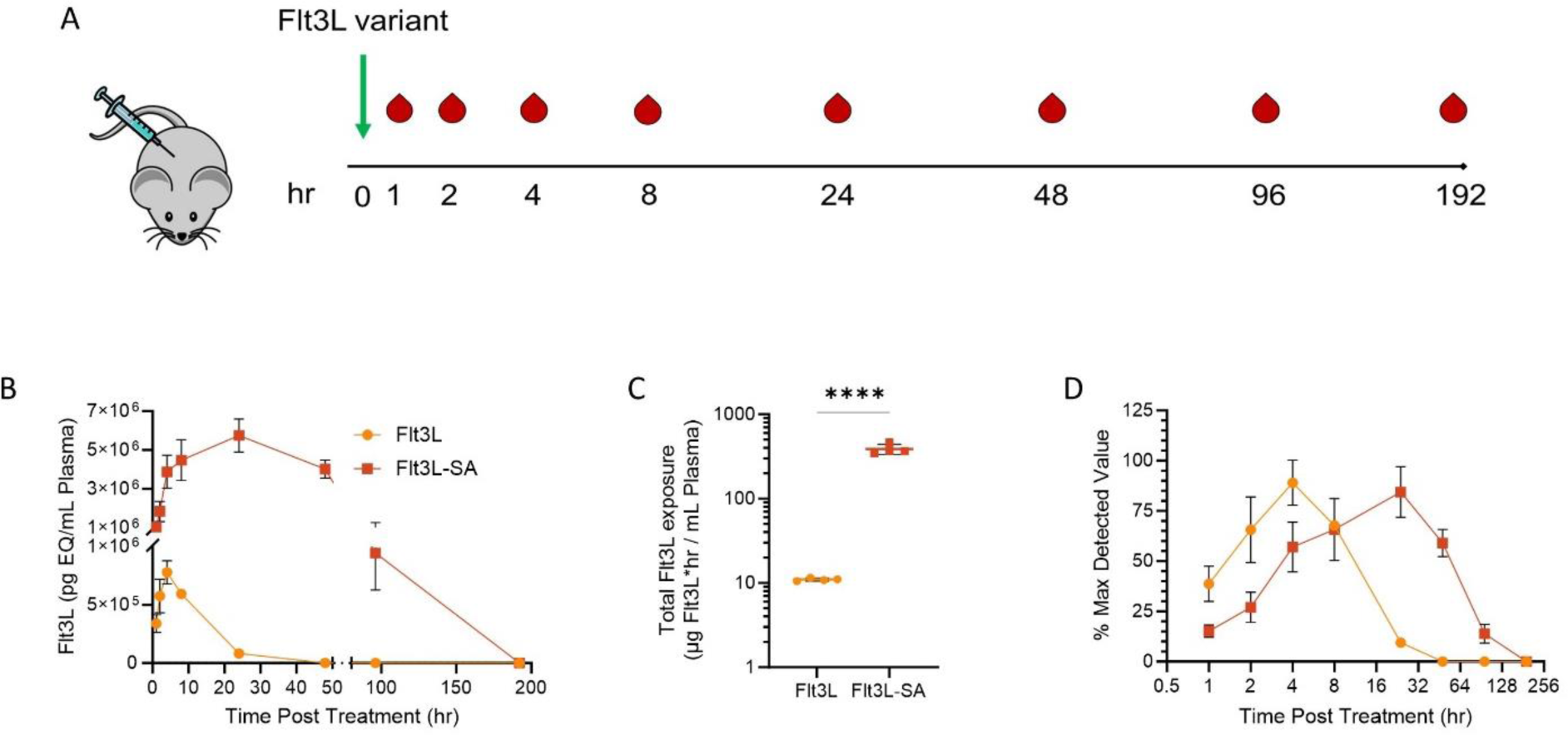
Serum albumin fusion significantly increases the circulatory half-life of Flt3L. (a) Schematic representation of the treatment and bleeding regimen. 10 µg of Flt3L or molar equivalent of Flt3L-SA was injected s.c. at time 0, with tail bleeds occurring at the indicated time points thereafter. (b) Flt3L concentration in the plasma at the indicated timepoints following s.c. injection with 10 µg Flt3L or molar equivalent of Flt3L-SA. (c) Total Flt3L exposure as denoted by the area under the curve demonstrated in (b). (e) Normalizing the values from (b) to the maximum detected value for each construct, Flt3L shows a rise to half maximum concentration of 1.3 hr and a fall back to half the maximum concentration of 10.6 hr. Flt3L-SA demonstrates an extended time to maximum concentration as well as extended half-life, equal to 4.1 and 55.8 hr, respectively. **** for p<0.0001.

### Pharmacodynamic properties of Flt3L-SA

Given that the Flt3L-SA fusion extended cytokine exposure *in vivo*, we next sought to determine the cellular changes following a single s.c. injection of Flt3L-SA. Mice received a single injection, and splenic immune populations were analyzed at the multiple times following the initial injection (Figure 3A). Peak splenic DC (CD11c^+^MHCII^+^) infiltration occurred 6 days following injection, reaching 8-fold higher numbers than those found in untreated mice (Figures 3B, Supplementary 1A, 2A). Interestingly, CD135 was undetectable on the DC population at 4 days following injection (Supplementary 1B, 2B), potentially indicative of continued cytokine signaling and receptor internalization,(67) as the Flt3L-SA is still present in the circulation at that time (Figure 2B). Within the DC subpopulations, the highest cell counts were observed 6 days after treatment in the cDC populations, including both cDC1 (IRF8^+^) and cDC2 (CD11b^+^).

**Figure 3:**
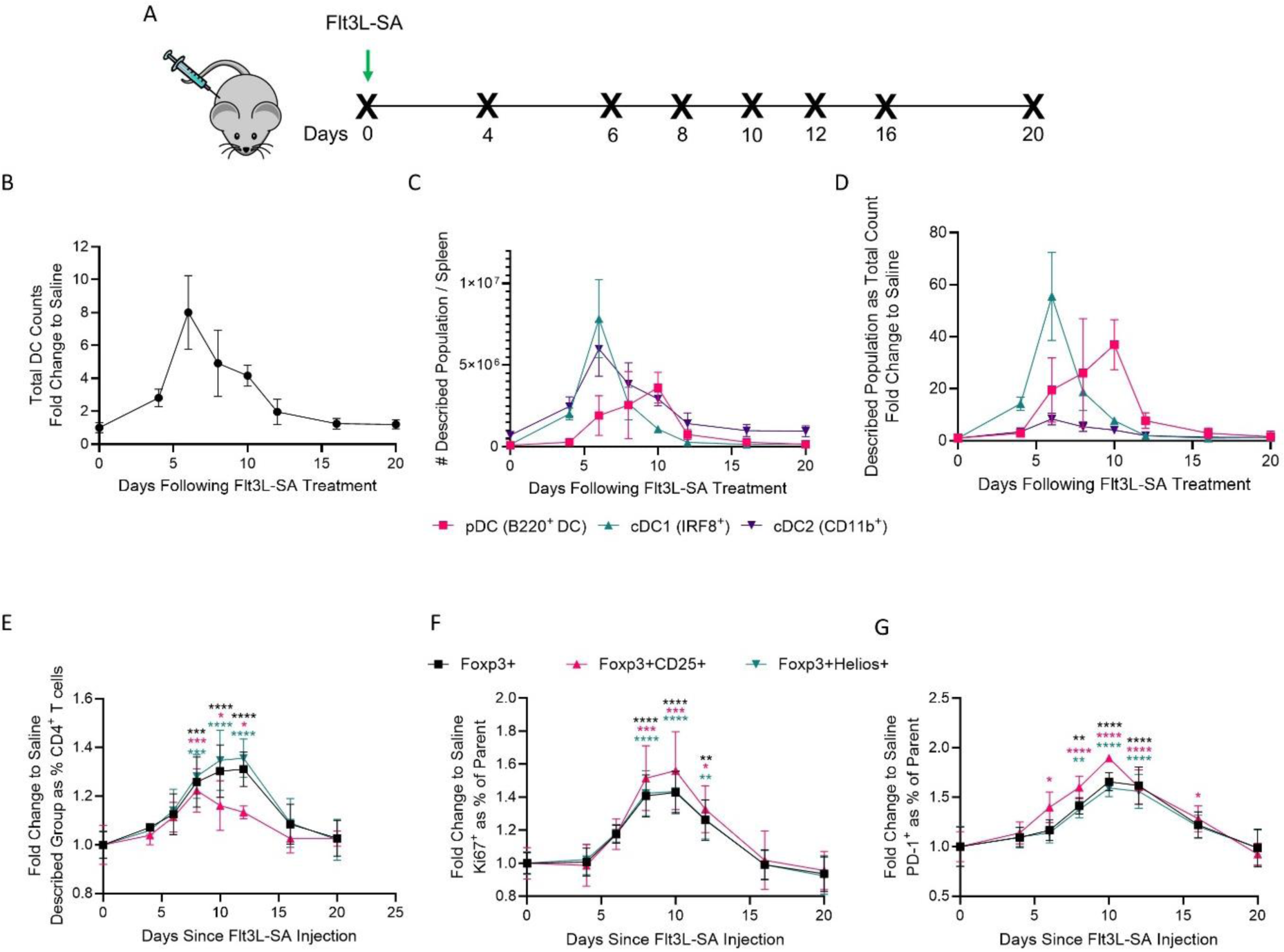
Pharmacodynamics of Flt3L-SA treatment demonstrate consistent expansion of DC and Treg populations after a single injection. (a) Treatment schematic demonstrating a single s.c. injection of 10 µg Flt3L-SA in the back of mice at time 0 with euthanasia and splenocyte isolation at times denoted with an X. (b) Fold change in total DC (CD11c^+^MHCII^+^) counts in the spleen as compared to saline-treated mice (time 0). (c) Number of pDC (B220^+^), cDC1 (IRF8^+^), and cDC2 (CD11b^+^) DC in the spleen at each indicated time. (d) Data indicated in (c) normalizing each population to time 0 to show relative changes to saline-treated mice. (e) Frequency of each described suppressive T cell population as a proportion of the total CD4^+^ T cell compartment after one Flt3L-SA injection, as normalized to time 0. Significance denotes each population as compared to time 0 via one-sided Students t-test. (f, g) Expression of markers of proliferation (Ki67) and (g) antigen recognition (PD-1) on cells from (e) as normalized to time 0. Significance denotes each population as compared to time 0 via one-way ANOVA with Dunnett’s multiple comparison. * for p<0.05, ** for p<0.01, *** for p<0.001, **** for p<0.0001.

Plasmacytoid DCs (pDC, B220^+^) also increased with a delay, peaking on day 10 (Figure 3C).(37, 38, 68–70) When normalized to untreated mice, it is notable that while cDC2s are typically the most abundant DC population, they exhibit the lowest relative increase with Flt3L-SA treatment. As such, cDC2s represent a stable percentage of the DC population (Supplementary 2C), whereas cDC1 and pDC populations showed expansion within the DC compartment, suggesting that DC progenitors experienced a pressure towards differentiating into these populations after Flt3L-SA treatment. Taken together, these data suggest that a single injection with Flt3L-SA expands the splenic DC compartment, with a bias towards cDC1 and plasmacytoid DC accumulation.

Though the precise mechanisms remain unknown, it has been previously established that Flt3L exposure leads to the expansion of Tregs.(48, 62, 64) Following the previous timeline (Figure 3A), we observed a significant increase in Foxp3^+^ and Foxp3^+^Helios^+^ T cells beginning 8 days after the initial injection, which peaked 10 to 12 days after treatment, before declining by day 16. The dynamics of Foxp3^+^ T cells displaying the high affinity IL-2 receptor, CD25, demonstrated a decreased peak response and a more gradual decline than the one exhibited by total Foxp3^+^ T cells (Figure 3E, Supplementary 1C). The aforementioned T cell subsets exhibited features denoting both proliferation and antigen exposure (Ki-67 and PD-1, respectively) suggesting activation and expansion of these T cell subsets (Figure 3F, G).

### Multi-dose regimen of Flt3L-SA leads to reduced activation state on DCs and Treg expansion

Previously published regimens involving Flt3L-induced DC proliferation involve daily treatment for 10 days;(37, 38, 50) therefore, we evaluated DC responses, focusing on DC accumulation and phenotype, following a treatment regimen that maintained Flt3L-SA exposure for that timeline (Figure 4A, Supplementary 3A-C). As expected, we observed a substantial expansion of DCs after Flt3L-SA treatment, but no major differences in DC populations between mice treated with an equimolar dose of WT Flt3L or saline control in this regimen (Figure 4B). Phenotypically, we observed decreased surface expression of activation markers MHCII and PD-L1 on DCs following Flt3L-SA treatment (Figure 4C). Furthermore, the total number of splenocytes expressing the TGF-β precursor LAP increased, including an increased LAP^+^ DC population (Figure 4D), comprised of both LAP^+^ cDC1 and cDC2 phenotypes (Figure 4E, Supplementary 3B). These results suggest that prolonged exposure to Flt3L-SA induces both a reduced activation state in the generated DCs as well as a potential pro-tolerogenic microenvironment, evidenced by LAP expression.(59, 71–74) However, when normalized to the parent DC population, we observed a decrease in the percentage of cDC1 and cDC2 expressing LAP, suggesting that Flt3L-SA may be expanding an already preexisting DC population, rather than inducing a specific cell state (Supplementary 4A-C). Additionally, we examined changes in the T cell compartment elicited by the sustained Flt3L-SA exposure. Though T cell proportions within the spleen fell with Flt3L-SA treatment, the absolute number of T cells rose, suggesting that the reduction in splenocyte proportion is most likely due to the observed DC expansion (Figure 4B, F). Notably, we observed a significant increase in both the frequency and number of Foxp3^+^CD25^+^ Tregs, indicating that prolonged treatment with Flt3L-SA can promote an expansion of this regulatory compartment (Figure 4G). These results together suggest that the Flt3L-SA fusion successfully reduces the burden of multiple injections and, under homeostatic conditions, may aid in enhancing a tolerogenic immune state.

**Figure 4:**
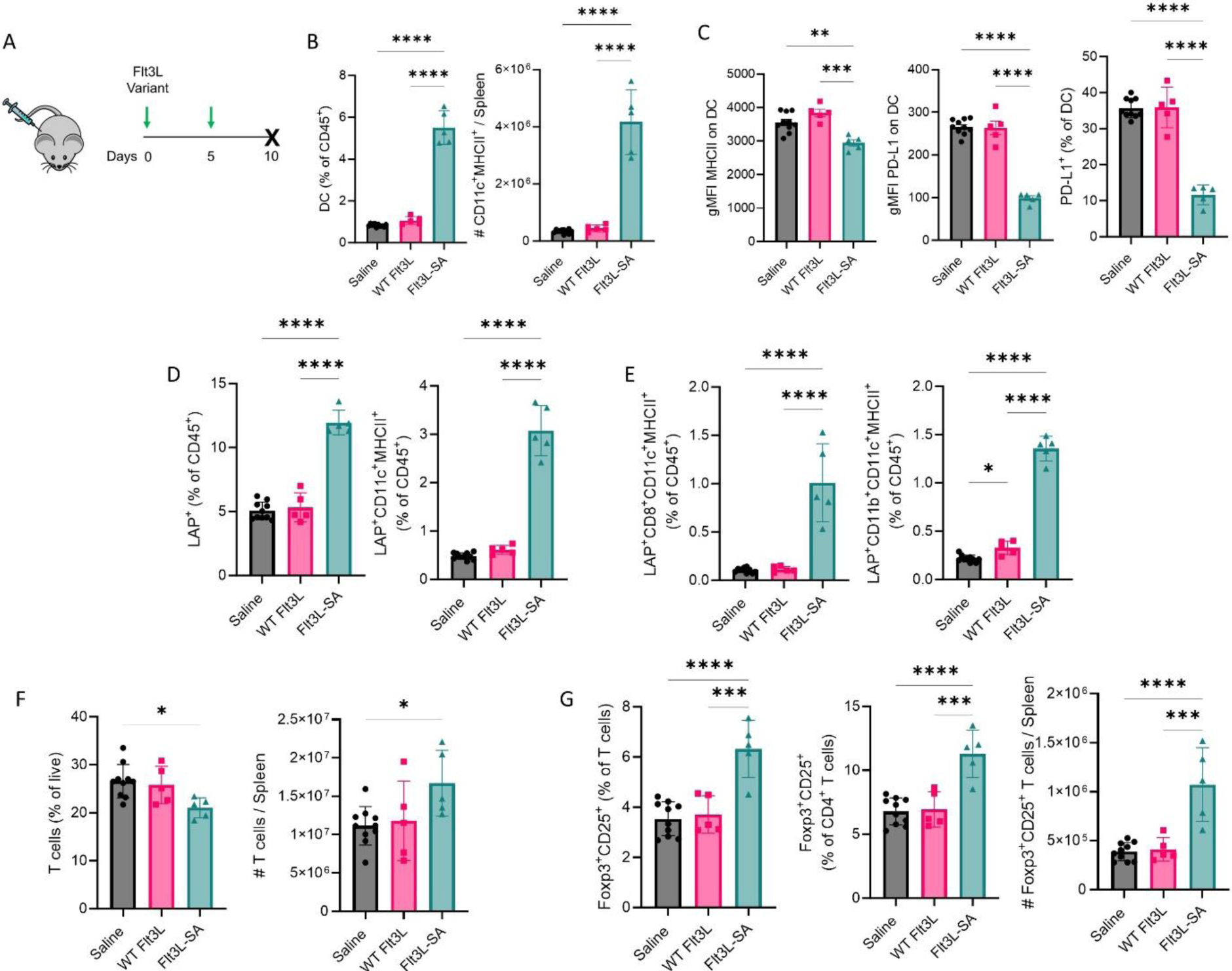
Flt3L-SA regimen expands non-inflammatory immune populations. (a) Schematic of treatment regimen. Mice were given 10 µg Flt3L or molar equivalent of Flt3L-SA s.c. on days 0 and 5 before euthanasia and splenocyte isolation on day 10. (b) DC (CD11c^+^MHCII^+^) cells as a proportion of total immune cells and absolute number based on total live counts in the spleen. (c) Phenotypic characterization of the cells in (b) for expression of markers of activation (MHCII and PD-L1). (d) Expression of the TGF-β precursor LAP in the spleen and proportion of hematopoietic cells expressing LAP and DC phenotypic markers (CD11c^+^MHCII+) via flow cytometry. (e) Expression of the TGF-β precursor LAP on cDC1 (CD8^+^) and cDC2 (CD11b^+^) DC subsets as proportion of total hematopoietic cells. (f) T cells (CD3e^+^) as portion of live cells in the spleen as well as absolute numbers in the spleen based on live count. (g) Treg (Foxp3^+^CD25^+^) characterization as a proportion of total T cells (left), CD4^+^ T cells (middle), and estimated count in the spleen (right). Error bars for gMFI graphs represent SEM, and are otherwise SD. Significance calculated using one-way ANOVA with Tukey’s multiple comparison correction. * for p<0.05, ** for p<0.01, *** for p<0.001, **** for p<0.0001.

### Flt3L-SA influences T cell primary education

We next examined how education of the CD4^+^ T cell compartment is affected during Flt3L-SA treatment, given the changes observed in the DC compartment. To do so, congenically-marked (CD45.1^+^) ovalbumin (OVA)-specific CD4^+^ T cells (OT-II) were adoptively transferred at the time of initial Flt3L-SA treatment, followed by intravenous (i.v.) OVA administration 3 days later, with both treatments repeated the next week (Figure 5A, Supplementary 5). Subsequent analysis of splenic immune populations revealed that in the bulk CD4^+^ T cell compartment, Flt3L-SA treatment expanded effector (CD44^+^CD62L^-^) CD4^+^ T cells (Figure 5B) and Foxp3^+^ CD4^+^ T cells, but decreased the population of cells expressing Gata3, a key transcription factor for TH2 fate (Figure 5C). Interestingly, analysis of the antigen-specific splenic OT-II cells, revealed that treatment with antigen did not influence the numbers or proportions of OT-II cells (Figure 5D). Unexpectedly, the effects of Flt3L-SA on the immune system diminished splenic CD4+ T cell education to the exogenous antigen OVA, whereas administration of Flt3L-SA with OVA reduced both Foxp3 and Gata3 induction in comparison to treatment with OVA alone (Figure 5E). These results suggest that TH2 and Treg education is impaired even in the presence of the expanded cDC1-dominated APC compartment, in the context of an intravenous antigen administration.

**Figure 5:**
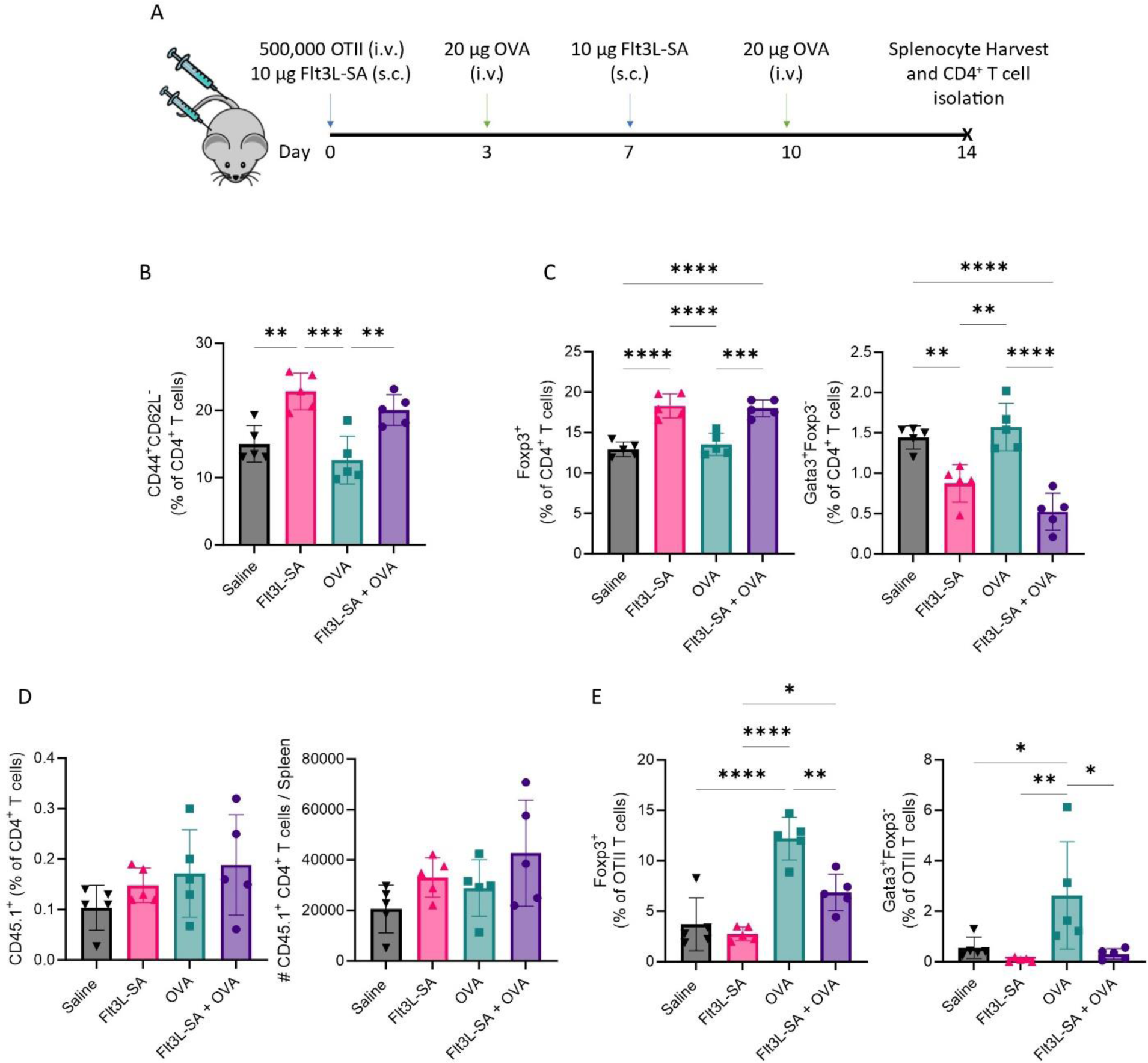
Antigen exposure during Flt3L-SA regimen modifies the elicited response and decreases CD4^+^ T cell effector differentiation. (a) Schematic representing the antigen exposure model with adoptive transfer of ovalbumin-specific CD4^+^ T cells (OTII) at the time of first s.c. Flt3L-SA treatment (day 0). Mice receiving Flt3L-SA were also treated on day 7, and mice getting antigen were given 20 μg ovalbumin (OVA) i.v. on days 3 and 10. Mice were euthanized and total CD4^+^ T cells were isolated from the spleen via magnetic sorting prior to flow cytometric staining. (b) Effector T cells (CD44^+^CD62L^-^) as a percentage of the total CD4^+^ T cell population. (c) Transcription factor staining related to T cell function including Treg (Foxp3^+^, left) and TH2 (Gata3^+^Foxp3^-^, right) as a percentage of total CD4^+^ T cells. (d) OVA-specific CD4^+^ T cells (CD45.1^+^ OTII) as a percentage of total CD4^+^ T cell population (left) and estimated number of cells in the spleen (right) as calculated as the cell population as a percent of live cells multiplied by the live splenocyte count. (e) Transcription factor staining related to T cell function including Treg (Foxp3^+^, left) and TH2 (Gata3^+^Foxp3^-^, right) as a percentage of OVA-specific CD4^+^ T cells. Error bars represent SD and significance is calculated using one-way ANOVA with Tukey’s multiple comparison correction. * for p<0.05, ** for p<0.01, *** for p<0.001, **** for p<0.0001.

### Flt3L-SA education reduces ADA development and lessens subsequent anaphylactic-like events

Considering the observed TH2 impairment and recent work tying TH2 phenotypes to high titer antibody formation in an enzyme-replacement setting,(75) we next sought to determine if Flt3L-SA treatment during the initial immune education period could reduce the development of ADAs against an exogenous enzyme drug. To this end, we developed a tolerance-induction therapeutic schedule using a low dose of an enzymatic drug, rasburicase, in combination with Flt3L-SA (Figure 6A). Rasburicase (Elitek™, Sanofi), is a highly immunogenic, homotetrameric uric acid oxidase enzyme, which is recombinantly produced from a fungal gene and has been approved by the U.S. FDA without PEGylation or other covalent modifications. Following this putative tolerance induction, we administered subsequent challenge doses of the drug at higher, therapeutically-relevant doses (Figure 6A). Analysis of plasma at endpoint revealed a significant decrease in total rasburicase-specific IgG in mice treated with this induction regimen, compared to mice in the challenge only group, i.e. proceeding directly to therapeutic doses (Figure 6B, Supplementary 6). The Flt3L-SA-mediated decrease in total IgG is potentially attributable to a decrease in antigen-specific IgG1, as this is the only isotype tested that did not also show a reduction in the low dose antigen group lacking Flt3L-SA co-administration (Figure 6C). IgG subclasses have been deemed relevant in anti-drug immune responses in that, in addition to basophil activation through IgE-mediated crosslinking, myeloid (primarily neutrophil) activation occurs through crosslinking of the low-affinity Fcγ receptor, FcγRIII (CD16), and IgG1 immune complexes are sufficient to cause hypersensitivity in mice and human patients.(76–78) Thus, reducing any and all IgG isotypes is likely to benefit patients in reducing hypersensitivity reactions to protein-based therapeutics.

**Figure 6:**
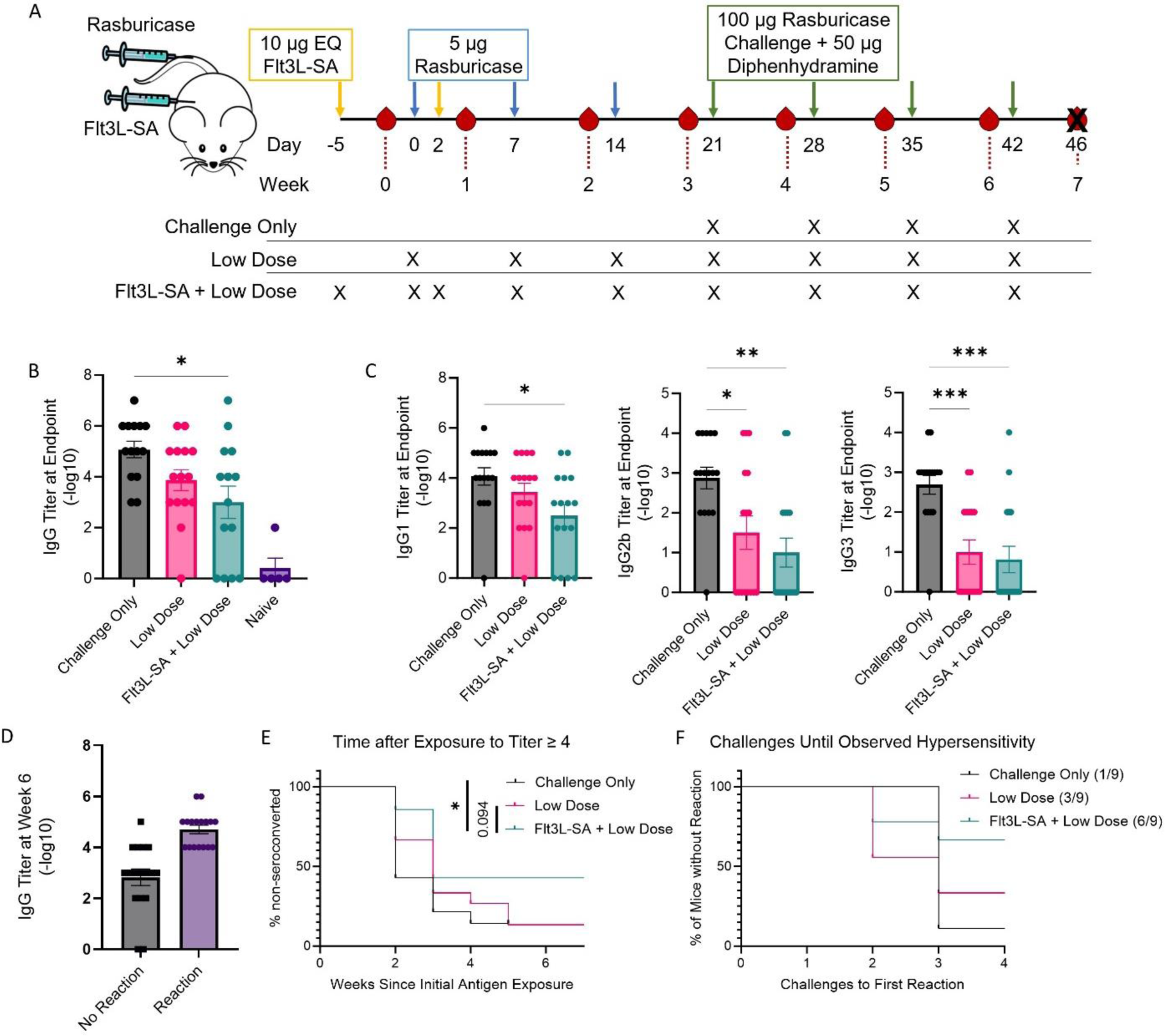
Flt3L-SA treatment in a tolerance induction regimen reduces anti-drug antibody levels and reduces pathogenic reactions to foreign enzymes. (a) Schematic representing the enzyme replacement challenge model with tolerance induction regimen. Treatments occur on the indicated days, with arrow color representing the treatment. Two induction doses with Flt3L or Flt3L-SA or saline were given with low doses of 5 µg rasburicase, followed by 5 therapeutic doses of 100 µg rasburicase. Flt3L-SA and diphenhydramine are both dosed s.c. and the rasburicase treatments are dosed i.v. Bleeds occur at the indicated weeks with the red drops highlighting the relative timing of the bleeds with respect to treatments. (b) Total rasburicase-specific IgG quantification via titration at experimental endpoint. Significance calculated by one-way ANOVA, significance against naïve not shown. N=5 for naïve group and n= 14-15 for the other three treatments. Error bars represent SEM. (c) Rasburicase-specific IgG subclass titration at experimental endpoint. Significance calculated by one-way ANOVA with n=14-15. Error bars represent SEM. (d) Mice grouped dependent on whether they demonstrated observable reactions at the final challenge against the measured IgG titer preceding the challenge dose. (e) Time until seroconversion to potentially anaphylactic levels of IgG (titer greater than 4) in plasma. Significance calculated via Kaplan-Meier testing. (f) Mice were evaluated 1 hr after each therapeutic challenge infusion for demonstrable anaphylaxis-like symptoms (hunched, ruffled, isolated, and/or cold). Censoring occurred at the first sign of reaction. Numbers indicate mice without demonstrable anti-drug infusion reactions after final challenge over total mice. * for p<0.05, ** for p<0.01, *** for p<0.001.

On the final, high-dose (i.e. therapeutic level) challenge, we stratified mice into those that developed observable anaphylactic-like reactions (including hunching, ruffling, isolation, and palpable temperature drop) and those that did not. Based on plasma, rasburicase-specific IgG titers preceding the challenge, we determined that seroconversion to a likely-anaphylactic state corresponded to a titer greater than or equal to 4 (Figure 6D). Using this criterion to segregate the mice into those likely or unlikely to react, we observed a significant benefit of the addition of Flt3L-SA to the low dose induction regimen as compared to no tolerance-induction, and a near-significant trend compared to low dose antigen alone (Figure 6E). Furthermore, in alignment with serology data, we noted a benefit in the number of mice that displayed an observable reaction following challenge when low dose induction occurred in combination with Flt3L-SA treatment (Figure 6F), showing that even a modest reduction in antibody titer can have a beneficial effect on symptomology. Altogether, these data suggest that addition of Flt3L-SA to an induction regimen is sufficiently capable of reducing the anti-drug response to lessen the risk of anaphylactic-like hypersensitivity reactions upon subsequent administration.

### Low dose antigen initiates a defective B cell response, which is amplified with the addition of Flt3L-SA

Given how the tolerance-inducing regimen reduced the ADA response, we next analyzed the splenic adaptive immune components at endpoint to probe how the B cell compartment was impacted by the induction therapy. Overall, we observed no differences in the bulk B cell compartment (Figure 7A, Supplementary 7A); however, all induction schemata reduced the prevalence of germinal center (GC) B cells (CD38^-^GL7^+^), indicative of a reduced affinity-maturation reaction (Figure 7B). Using avidin-based antigen tetramerization and fluorescent labeling, we analyzed rasburicase-specific B cells via their surface-bound B cell receptor (Figure 7C), in which we observed a significant decrease in the proportion of B cells specific to rasburicase in both induction schema and a near-significant decrease in the number of these cells when Flt3L-SA was added (Figure 7D). We further characterized the rasburicase-specific B cells via surface expression of CD80 as a marker of activation and T cell cross-talk. We observed that the tolerance-inducing regimens involving low-dose rasburicase significantly decreased CD80 expression on antigen-specific B cells when compared to those isolated from mice that were not tolerized. However, only the group that treated with Flt3L-SA in the induction regimen showed no significant difference in CD80 expression between the antigen-specific and the bulk B cell compartments (Figure 7E), which may be indicative of a disruption in B-to-T cell communication.(79) Similar to the observations of the bulk GC B cell compartment, both induction schema decreased antigen-specific GC B cells (Figure 7F). Additionally, we observed tolerization-induced reductions in antigen-specific memory B cells (CD38^+^IgD^-^), with a trending decrease in total number of these cells only in the group that received Flt3L-SA (Figure 7G).

**Figure 7:**
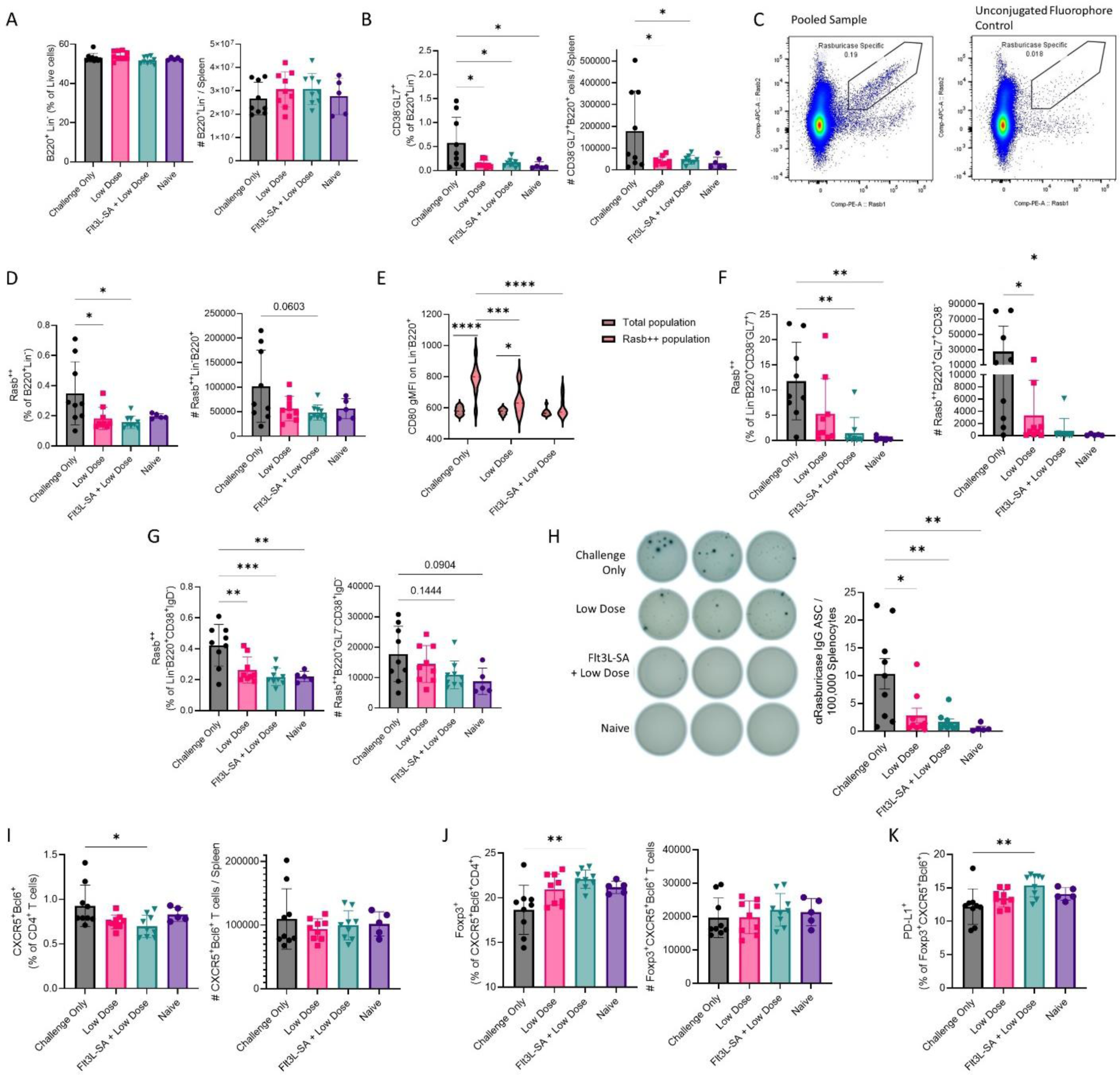
Flt3L-SA tolerance regimen decreases the antigen-specific B cell compartment and alters B and T cell communication networks. (a) At endpoint of experiment in Figure 5, B cell (B220^+^Lin^-^) enumeration as percent of live splenocytes and estimation of total B cells in the spleen. (b) Germinal center (GC) B cells (CD38^-^GL7^+^) as percent of total B cells and estimated number in the spleen. (c) Representative gating for antigen-specific B cells using fluorescent rasburicase probes vs. unconjugated fluorophore control. (d) Quantification of antigen-specific B cells (double positive for APC and PE conjugated probes) as a percent of total B cells as well as estimated numbers in the spleen. (e) Comparative characterization of expression of the costimulatory marker CD80 on total B cells as well as antigen-specific B cells in (d). Significance calculated using RM two-way ANOVA with Sidak’s multiple comparison correction with a single pooled variance. (f) Quantification of antigen specific GC B cells as a percent of total GC B cells as well as estimated total number of antigen specific GC B cells in the spleen. (g) Quantification of antigen-specific memory B cells (CD38^+^ IgD^-^). Antigen-specific cells as a percent of total memory B cells (left) and estimated number of antigen-specific memory B cells in the spleen (right). (h) Quantification of antigen specific IgG secreting cells via ELISPOT capturing all IgG and detecting with biotinylated rasburicase. Representative wells for one mouse for each condition (left) and total enumeration (right). Each point represents one mouse, the average of three wells with error bars for SEM. Unless otherwise specified, each point represents one mouse with error bars for SD and significance calculated with one-way ANOVA with Tukey’s multiple comparison. (i) Percent of CD4^+^ T cells with a Tfh phenotype (CXCR5^+^Bcl6^+^) as well as estimated number in the spleen. (j) Quantification and phenotype of T follicular regulatory (Tfr) cells. Percent of cells in (a) expressing Foxp3 (left) and estimated number of Tfr cells in the spleen (middle). (k) Percent of Tfr expressing PD-L1 thus capable of suppressing lymphocyte reactions. Each data point represents one mouse with error bars for SD. Statistics calculated via one-way ANOVA between all groups with Tukey’s multiple comparison correction. * for p<0.05, ** for p<0.01, *** for p<0.001, **** for p<0.0001.

Finally, we sought to directly analyze the IgG-secreting compartment via antigen-specific ELISpot, wherein we noted reductions in anti-rasburicase IgG-secreting cells in mice given induction doses of rasburicase (Figure 7H). These data suggest a potential role of Flt3L-SA inducing a defect in B cell maturation pathways, especially when considering the reduction in antigen-specific GC B cells and memory populations.

### Flt3L-SA induced tolerance skews follicular T cells towards a regulatory fate

Considering how Flt3L-SA treatment increases systemic Treg presence, we next examined bulk T cell readouts of tolerance. We found that Flt3L-SA tolerization significantly decreased the frequency of T follicular helper cells (Tfh, CXCR5^+^Bcl6^+^) in the spleen (Figure 7I, Supplementary 7B, C). Additionally, we observed that of these cells, only Flt3L-SA-induced tolerance led to a significantly increased percentage of Tfh cells co-expressing Foxp3, a subset known as T follicular regulatory cells (Tfr) (Figure 7J), as well as trends towards increased bulk Tregs (Supplementary 8A, B). Indeed, DCs have been shown to be essential in priming and initiating the Tfr differentiation process.(80, 81) We also noted a significant increase in PD-L1 expressing Tfr cells only in the group tolerized during Flt3L-SA treatment (Figure 7K), which aligns with recent work on direct Tfr inhibition of T and B cell reactions in germinal centers.(82, 83) These results support the hypothesis that immune education under the influence of Flt3L-SA induces abrogation of inflammatory T cell education and increases regulatory signaling to control immune reactions.

## Discussion

In this study, we developed and characterized the potential of Flt3L fused to serum albumin to induce tolerance at initial antigen exposure. Fusion to SA extends the half-life of Flt3L by roughly 5-fold and increases the overall exposure to the cytokine by a factor of 30, similar to pharmacokinetic profiles in other work involving Flt3L fusions.(40, 61) Whereas most recent research related to Flt3L has emphasized the utility of the protein in enhancing a cross-presenting DC compartment for vaccination and anti-tumor immunity in the context of mounting a pro-inflammatory immune response,(37–40, 50) we introduce a different strategy emphasizing the tolerance mechanisms of DCs at play in the absence of inflammatory signals. With the addition of low dose exogenous antigen in the absence of immunomodulating agents, such as adjuvants or immunosuppressants, Flt3L-SA-expanded DCs were capable of changing how the adaptive immune system reacted to administered foreign protein. Specifically in this regimen, de novo education of antigen-specific OTII cells towards Tregs and Gata3^+^ phenotypes were reduced when foreign antigen was given in the context of Flt3L-SA expanded DCs, even though bulk Tregs had been expanded as expected.

Evidence of an abrogated adaptive response further led us to explore disruption of antigen recognition to a protein therapeutic upon initial exposure. To study the effect of Flt3L-SA on adaptive immune responses, we developed an induction regimen in which tolerance to a known immunogenic protein, rasburicase, may be established and measured over time.

Rasburicase (Elitek™) is an FDA approved, homotetrameric enzyme used to convert uric acid to a more soluble metabolite, allantoin, but due to its immunogenicity is not approved for long-term use.(84, 85) When mice were given an induction regimen of Flt3L-SA and low-dose rasburicase and then crossed-over to therapeutic (challenge) doses of rasburicase (in the absence of Flt3L-SA), we observed a significant reduction in systemic IgG antibodies, including the IgG1 subclass, related to type 2 immune responses.(86) Furthermore, it has been shown that a reduction in antibodies against uricase enzymes is therapeutically beneficial in maintaining enzyme function in human patients.(87–90) Finally, we also noted a significant reduction in the risk of mice developing anaphylaxis-like symptoms upon infusion when tolerization occurred during the Flt3L-SA treatment regimen. This effect suggests promise that the reduction in antibodies could benefit patients by preventing immune reactions to infused drugs when used upon first exposure.

These studies are not without their limitations. Future work in this project will involve determining whether the reduced antibody reaction is beneficial in maintaining enzyme functionality. Unlike humans, mice express an endogenous uric acid oxidase enzyme which makes assays to probe the functionality or clearance of the enzyme in vivo are not trivial. However, it has been shown in humans that a slight reduction in antibody titers is sufficient to provide benefit in patients treated with pegloticase, a less immunogenic (due to PEGylation and mammalian origin) recombinant uric acid oxidase treatment for refractory gout.(87) Additionally, we would like to probe antigen-specific T cells in the rasburicase treated mouse model to determine how such a compartment is changing in response to treatment with Flt3L-SA and compare results to the OVA/OT-II system. Due to the enzymatic function and homotetrameric structure of rasburicase, it may be interesting to assess how these aspects affect T cell education, e.g. in contrast to immune responses to a monomeric protein drug. Finally, our focus here is on the use of Flt3L-SA with low-dose drug as an induction regimen in treatment-naïve subjects, i.e. just prior to the time of initial drug exposure.

Overall, concurrent with the clinical growth of biologic therapeutics, inflammatory anti-drug reactions will continue to be a problem in maintaining efficacy of the biologic agents, including those that have been engineered using non-homologous regions.(91–95) One arm seeking to reduce immunogenicity to these drugs is the research into therapeutic-intrinsic factors. Direct chemical conjugation to the drug may shield immune epitopes to passivate the drug(96) or may actively direct tolerogenic uptake of the therapy.(13, 14, 16, 97) Alternatively, amino acid sequence mutations following *in silico* immunogenicity prediction have been employed to reduce immune recognition(98–102) or to insert epitopes to direct Treg activation and development.(103, 104) Another arm in the battle against anti-drug reactions is that of co-therapies given in course with the biologic drug to change the microenvironment in which the immune system encounters the treatment. These mostly include the use of immunomodulating agents such as methotrexate to prevent or reduce the prevalence of anti-drug reactions(19, 105–109) but may come with the additional side-effect of broad immunosuppression. One engineering tactic in particular, known as ImmTOR nanoparticles, uses encapsulated rapamycin to direct the immunosuppressive drug towards antigen-presenting cells concurrent with early administration of the biologic agent.(110) This strategy promotes generation of stable, antigen-specific tolerance while mitigating the broad immunosuppression of free rapamycin. Herein, we demonstrate the utility of an antigen-agnostic regimen that employs soluble recombinant proteins (in the absence of broad immunosuppressants), with their relative ease of clinical translation, and focuses on changing the context of the immune system, with an emphasis on the DC compartment, as an induction regimen to create a tolerance-promoting state during initial drug exposure. This tolerance-induction regimen reduced the anti-drug antibody burden and the associated complications after repeated dosing at therapeutic levels.

## Materials and Methods

### Sex as a biological variable

All mouse experiments were performed using female mice to allow for cohousing of animals without risk of incidental inflammation due to aggression and consistent adoptive cell transfer across groups.

### Animals

Wild type mice were purchased from Jackson Labs, and congenically marked F1 CD45.1 OTII mice were bred in-house and housed at the University of Chicago Animal Facility. Mice were randomized with every experiment to ensure distribution of cage affects across all groups.

Injections were diluted to the appropriate concentration in 50 µL sterile saline for s.c. injections or 100 µL for i.v. or i.p. treatments. Diphenhydramine treatments occurred s.c. into the flank of the mouse 45 min prior to challenge. Mice were euthanized via CO_2_ asphyxiation as approved by UChicago ARC.

### Protein production and purification

Plasmids encoding Flt3L or Flt3L-SA on a pcdna3.1 vector, made in house, were transfected into HEK293F cells using linear, 25kDa polyethyleneimine as previously described.(111) Transfected cells were then incubated at 37°C for one week with constant shaking before protein harvesting. At harvest, cells were removed by centrifugation at 4000G for 10 min before sterile filtration and pH adjustment. Supernatant was then loaded over a HisTrap HP (Cytiva) column on a GE AKTA Avant FPLC, and the protein was washed and eluted with varying amounts of imidazole. Protein eluted was further purified over a superdex 200pg column (Cytiva) in PBS. Protein concentration was determined via absorbance at 280 nm via NanoDrop (ThermoScientific) using the predicted molecular weight and extinction coefficient. The final protein was then tested for purity via SDS-PAGE using lamelli buffer for non-reducing conditions and the same buffer with added 0.14 M β-mercaptoethanol for reducing conditions. LPS testing was performed using a HEK TLR4 cell line with quantiblue to confirm endotoxin free samples before aliquoting and freezing.

### CD135 binding ELISA

High-bind plates (Corning 9018) were coated by incubating overnight at 4°C with 10 nM recombinant CD135 (R&D 768-F3-050) in 50 nM sodium bicarbonate buffer at pH 9.2. Following coating, plates were washed in PBST before blocking in 1x reagent diluent (R&D DY995) for 2 hr at RT. Samples were then diluted to the indicated concentration in triplicate using the same reagent diluent. Blocked wells were washed and then sample was added and incubated for 1 hr. Samples were removed before addition of the detection antibody (R&D 841481) at the recommended concentration and incubated for 1 hr followed by washing and addition of Streptavidin-HRP (R&D 890803) at the indicated concentration for a 30 min incubation. Wells were then washed before the addition of TMB (Millipore ES-001). TMB was incubated while protected from light until saturation was reached, at which point the reaction was stopped using 10% sulfuric acid. Quantification occurred by reading the absorbance at 450 nm and 570 nm. Data was then plotted as the absorbance at 450 nm minus absorbance at 570 nm.

### BMDC generation

BMDC were generated according to a modified Lutz protocol.(112, 113) Briefly, bone marrow was flushed from the long bones of healthy 6–15 week-old C57BL/6 mice into RPMI 1640 media and filtered over 100 µm filters. On day 0, 3 million nucleated cells were plated in 10 mL of a modified Lutz media (RPMI 1640 supplemented with 10% FBS, 1% Pen/Strep/L-glutamine, 50 µM β-mercaptoethanol, 25 mM HEPES, 20 ng/mL GM-CSF, 200 ng/mL Flt3L) in 100 mm non-tissue culture treated petri dishes. Cells were fed by the addition of 10 mL of the initial media (with GM-CSF and Flt3L) on day 3. On day 6, media was refreshed by removing 10 mL and centrifuging the cells before resuspension in 10 mL of fresh, complete media (GM-CSF and Flt3L). Non-adherent cells were harvested on day 9 for use and starved for 2 hr in incomplete RPMI 1640.

### Phospho-ERK1/2 Assays

Starved Flt3L-generated BMDCs were plated (1 million cells for western blots, 500,000 cells for phospho-flow) in 100 µL of incomplete RPMI. For Western-based assays, 100 µL of 4 µM Flt3L variant in incomplete RPMI was added to each tube for the indicated time before dilution with 1 mL of ice-cold PBS and centrifugation. Pelleted cells were then lysed using RIPA buffer (Thermo 89901) with protease and phosphatase inhibitor added (Thermo A32959, 1 tablet/10 mL RIPA). Cells were incubated in the RIPA buffer for 5 min on ice before centrifugation at 17,000 G for 10 min. Supernatant was taken and total protein quantified via BCA (Thermo 23227). 8 µg of protein was then separated via SDS-PAGE before transfer to a PVDF membrane. Blocking occurred using 5% BSA in TBST and antibody probing using 2% BSA in TBST at the indicated dilutions: α-pERK1/2 (Biolegend 369502) 5000x, HRP-α-Mouse (CST 7076S) 10,000x, α-total ERK1/2 (Biolegend 686902) 5000x, HRP-α-Rat (Jackson Immuno 112-036-003) 10,000x. The membrane was stripped between phospho-ERK1/2 blotting and total ERK1/2 blotting using Restore Stripping Buffer (Thermo 21059). For phospho-flow assays, warmed cytokine stimulations in incomplete RPMI were added to each well before incubation for exactly 5 min. At the end of the incubation period, 50 µL of warmed 5x Lyse/Fix buffer (BD 558049) was added to each well before incubation at 37°C for 10 min. Cells were then washed with PBS before resuspension in ice-cold Perm Buffer (BD 558050) and a 15-min incubation on ice. Cells were then washed twice in FACS buffer (PBS + 2% FBS + 1mM EDTA) before staining with anti-pERK1/2 (Biolegend 369506 100x, as described in methods for flow cytometry staining) and acquisition on a BD Fortessa.

### Blood and plasma collection

Blood was collected via tail vein puncture (for pharmacokinetic studies) or submandibular venous puncture (for antibody titrations) into Lithium-Heparin coated tubes (Starstedt). For plasma collection, tubes were then centrifuged at 10,000G for 10 min before collection of the plasma and subsequent freezing into PCR tubes.

### Pharmacokinetics study

Once plasma for all timepoints had been collected as described, Flt3L content in the plasma was quantified via ELISA (R&D Dy427) using Flt3L standard for mice treated with WT Flt3L and using an equimolar Flt3L-SA standard for Flt3L-SA treated mice. Plasma was diluted 100-10,000x and the lowest dilution for each timepoint which did not oversaturate the standard was taken and converted before accounting for dilution.

### Preparation of single-cell suspensions

After euthanasia, spleens were harvested into 0.5 mL of complete DMEM and placed on ice. Once spleens from all mice were removed, they were pushed through a 70 µm filter and washed with incomplete DMEM. Suspensions were then centrifuged at 1750 RPM for 7 min before resuspension in 3 mL of ACK lysis buffer (Thermo 1049201) followed by a 5 min incubation before dilution in incomplete DMEM. Cells were then centrifuged as previously stated before counting and a final resuspension in complete DMEM at a concentration of 20 million cells/mL.

### OT-II isolation and adoptive transfer

Splenocytes were isolated as previously described and red blood cells lysed before CD4^+^ T cell isolation using the STEMCell isolation kit (19852) according to manufacturer instructions. Isolated cells were then resuspended to the desired number of cells in 100 µL of sterile, incomplete DMEM for intravenous injection via tail vein.

### Protein transit inhibition and flow staining

2 million isolated cells were plated per sample before washing with plain PBS. Viability stain was diluted in plain PBS at indicated concentration with the addition of Fc Block before adding 50 µL/well and incubation on ice for 15 min. Viability dye was quenched by washing with FACS buffer (PBS + 2% FBS + 1 mM EDTA) before addition of surface staining antibodies in a 1:1 dilution of Brilliant Stain Buffer (BD 563794) in FACS buffer. Surface staining occurred for 20 min at RT before washing in plain PBS and subsequent fixation. Samples not requiring intracellular staining were fixed for 20 min on ice using 2% PFA in PBS. Samples requiring only intracellular cytokine staining were fixed for 20 min on ice using the BD Cytofix/Cytoperm kit (BD 554714) before washing and intracellular staining for at least 1 hr at 4°C. Samples staining for nuclear factors (with or without cytokine staining) were fixed using the Foxp3 Transcription factor staining set (Thermo 00-5523-00) for 45-60 min on ice before intracellular staining for at least 1 hr at 4°C. Conditions where LAP staining was performed had a pre-incubation step in GolgiPlug (BD 555029) and GolgiStop (BD 554724) to increase sensitivity of the staining. Briefly, 2 million isolated cells were plated in Complete RPMI (RPMI + 10% FBS + 1% P/S) with 1x each inhibitor according to manufacturer instructions. Cells were then incubated at 37°C for 4 hr before proceeding with flow staining, using the CytoFix/CytoPerm (BD 554714) kit for intracellular staining. After fixation and intracellular staining, cells were washed and resuspended in FACS buffer for data acquisition on a 5 laser BD fortessa or 5 laser Cytek aurora.

### Anti-rasburicase titering ELISA

High-bind plates (Corning 9018) were coated by incubating overnight at 4°C with 1 µg/mL (for total IgG) or 10 µg/mL (for IgG1, IgG2b, and IgG3) rasburicase in 50 nM sodium bicarbonate buffer at pH 9.2. Following coating, plates were washed in PBST before blocking in 1x casein buffer (Thermo) for 2 hr at RT. Plasma was then serially diluted into the blocking buffer in 10-fold dilutions beginning at 100x (titer of 2) and finishing at a titer of 9. Blocked wells were washed and then sample was added and incubated for 1 hr. Sample was removed before addition of HRP-conjugated α-mouse IgG, α-mouse IgG1, α-mouse IgG2b, α-mouse IgG3, (all from Southern Biotech) diluted 5000x in blocking buffer and incubated for 1 hr. Wells were then washed before the addition of TMB (Millipore ES-001). TMB was incubated while protected from light for exactly 18 min, at which point the reaction was stopped using 1% hydrochloric acid + 3% sulfuric acid. Quantification occurred by reading the absorbance at 450 nm and 570 nm. To quantify a titer as positive, each well had the 570 nm reading subtracted from the 450 nm reading. Following this, background absorbance and standard deviation thereof was calculated using at least 3 blank wells for each plate. Test wells then had average background subtracted as well as 4x the standard deviation. Any well with a value greater than 0.01 was considered a positive titer, and the final positive titer was reported. Any mice in which the titer of 2 showed no signal were reported as a titer of 0. AUC was calculated as the Riemann sum of the titer vs background-subtracted absorbance curve for each time-point.

### Fluorescent rasburicase probe production

Rasburicase was biotinylated using EZ-link NHS-biotin (Thermo Scientific). Unreacted NHS-biotin was removed using Zeba spin desalting columns, 7 kDa MWCO (Thermo Scientific). The extent of biotinylation was measured using the QuantTag Biotin Quantification kit (Vector Laboratories) to ensure 1:1 molar ratio of rasburicase and biotin. Biotinylated rasburicase was reacted for 20 min on ice with 4:1 molar ratio of biotin to streptavidin-conjugated PE or streptavidin-conjugated APC (Biolegend). Streptavidin-conjugated FITC (BioLegend) was reacted with excess free biotin to form a non-antigen-specific streptavidin probe as a control. Cells were stained for flow cytometry with all three streptavidin probes at the same time as other fluorescent surface markers at a volumetric ratio of 1:400 for the PE and APC probes, and 1:100 for the FITC probe.

### Anti-rasburicase ASC ELISPOT

Plates (Millipore MAIPSWU) were activated using 70% EtOH for exactly 2 min before 3 washes with sterile PBS. Anti-mouse IgG capture antibody (Mabtech) diluted to 15 µg/mL in PBS was then added to each well before incubation overnight at 4°C. The next morning, plates were washed with complete media then and incubated at 37°C for at least 2 hr to block. Plates were then washed before adding 300,000 or 150,000 splenocytes/well in triplicate (6 wells/mouse, 3/condition). Plates were incubated at 37°C for 18 hr without movement or jostling. After the incubation, plates were washed with PBS and biotinylated-rasburicase (as prepared for the fluorescent antigen probes) was added at 1 µg/mL of PBS + 0.5% BSA (w/v) and incubated for 2 hr at RT. Plates were then washed with PBS before the addition of 1x streptavidin (Biolegend) in PBS + 0.5% BSA (w/v) and incubation for 1 hr at RT. Finally, plates were washed before the addition of TMB (Mabtech) and incubated until spots were visible (10 min) followed by quenching the reaction by washing with DI water. Plates were then dried in the dark before imaging using a CTL ImmunoSpot Analyzer to image, count spots, and perform quality control.

### Antibodies and Stains Used

All purchased flow cytometry stains can be found in Supplemental Table 1.

## Data Analysis

All data was plotted and analyzed on GraphPad Prism v10 software using statistic tests noted in the figure legends. Flow cytometric data was analyzed using FlowJo v10.

## Study Approval

All procedures were approved by the University of Chicago Institutional Animal Care and Use Committee and performed on protocol 72449.

## Data Availability

All data needed to evaluate the conclusions in the paper are present in the paper itself or included in the Supplementary Materials. Values for all data points in graphs are reported in the Supporting Data Values file.

## Author Contributions

Conceptualization, A.T.A., R.P.W., and J.A.H.; Methodology, A.T.A., R.P.W., S.G., L.A.H., S.C., J.E.G.M., and J.A.H.; Investigation, A.T.A., R.P.W., K.C.R., S.G., A.S., L.T.G., A.J.S., A.L.L., L.A.H., S.C., J.E.G.M., L.G.R., and J.A.H.; Visualization, A.T.A., S.G., and J.A.H.; Writing – Original Draft, A.T.A. and J.A.H.; Writing – Reviewing and Editing, A.T.A., R.P.W., K.C.R., A.J.S., and J.A.H.; Funding Acquisition, J.A.H.; Supervision, J.A.H.

## Acknowledgements

We thank the Cytometry and Antibody Core Facility at the University of Chicago (RRID: SCR_017760). We thank the UChicago Animal Resources Center (RRID:SCR_021806). We thank The University of Chicago HIM Facility (RRID:SCR_017916). We thank the Comprehensive Cancer Center DNA Sequencing Facility for sequencing all vectors. We thank Yue Wang for her laboratory support throughout this project.

