## Supplemental Data and Table for "Engineered Flt3L Drives Tolerogenic State to Attenuate Anti-drug Antibody Responses"

**Supplementary Table 1: Antibodies and stains used for flow cytometry**

| Antigen | Fluorophore | Vendor | Catalog # | Staining dilution |
| --- | --- | --- | --- | --- |
| Phosphor-ERK1/2 | PE | Biolegend | 369506 | 1:100 |
| CD19 | FITC | Biolegend | 152404 | 1:200 |
| CD11c | BV605 | Biolegend | 117333 | 1:200 |
| I/A-I/E | PerCP-Cy5.5 | Biolegend | 107626 | 1:200 |
| B220 | BUV496 | BD Biosciences | 612950 | 1:200 |
| PDCA1 | BV510 | BD Biosciences | 747607 | 1:200 |
| IRF8 | APC | Invitrogen | 17-9852-82 | 1:200 |
| CD11b | BV785  PE-Cy7 | Biolegend  Biolegend | 101243  101216 | 1:200  1:200 |
| CD135 | BV421 | BD Biosciences | 562898 | 1:100 |
| CD3ε | BUV395  BV605  PerCP-Cy5.5 | BD Biosciences  Biolegend  Biolegend | 563565  100351  100328 | 1:200  1:800  1:400 |
| CD4 | BUV496  BV785 | BD Biosciences  BD Biosciences | 612952  740844 | 1:200-400  1:200 |
| Helios | AF488 | Biolegend | 137223 | 1:200 |
| Ki-67 | PE | Biolegend | 652404 | 1:200-400 |
| PD-1 | BV785 | Biolegend | 135225 | 1:200 |
| CD25 | BV605  APC-Cy7  BV650 | Biolegend  Biolegend  Biolegend | 102035  102026  102038 | 1:200  1:200  1:200 |
| Foxp3 | BV421  FITC  PE-Cy7 | Biolegend  BD Biosciences  Invitrogen | 126419  560403  25-5773-82 | 1:50  1:200  1:50 |
| CD45 | APC-Cy7  APC | Biolegend  Biolegend | 103116  103124 | 1:200  1:200 |
| CD8α | BUV737  BUV395 | BD Biosciences  BD Biosciences | 612759  563786 | 1:200  1:400 |
| LAP | BV421 | Biolegend | 141408 | 1:200 |
| PD-L1 | BV786 | BD Biosciences | 741014 | 1:200 |
| CD44 | BV421 | Biolegend | 103039 | 1:200 |
| CD62L | BUV805 | BD Biosciences | 741924 | 1:800 |
| Gata3 | AF488 | Invitrogen | 53-9966-42 | 1:50 |
| CD45.1 | AF647 | Biolegend | 110720 | 1:100 |
| IgD | BUV395 | BD Biosciences | 564274 | 1:200 |
| CD138 | BV605 | Biolegend | 142531 | 1:200 |
| CD80 | BV650 | Biolegend | 104732 | 1:50 |
| IgM | BV786 | BD Biosciences | 743328 | 1:50 |
| Dump gate –  F4/80  CD11c  Gr-1  CD4  CD8  Streptavidin | FITC | Biolegend  Biolegend  Biolegend  Biolegend  Biolegend  Biolegend | 123108  117306  108406  116003  100706  405202 | 1:100  1:100  1:100  1:100  1:100  1:100 |
| GL-7 | PerCP-Cy5.5 | Biolegend | 144610 | 1:200 |
| CD38 | APC-Cy7 | Biolegend | 102727 | 1:200 |
| CXCR5 | BV421 | Biolegend | 145512 | 1:100 |
| Bcl6 | PE | BD Biosciences | 561522 | 1:100 |
| Viability | eFluor780  eFluor455  ZombieAqua | Invitrogen  Invitrogen  Biolegend | 65-0865-14  65-0868-14  423101 | 1:500  1:500  1:800 |

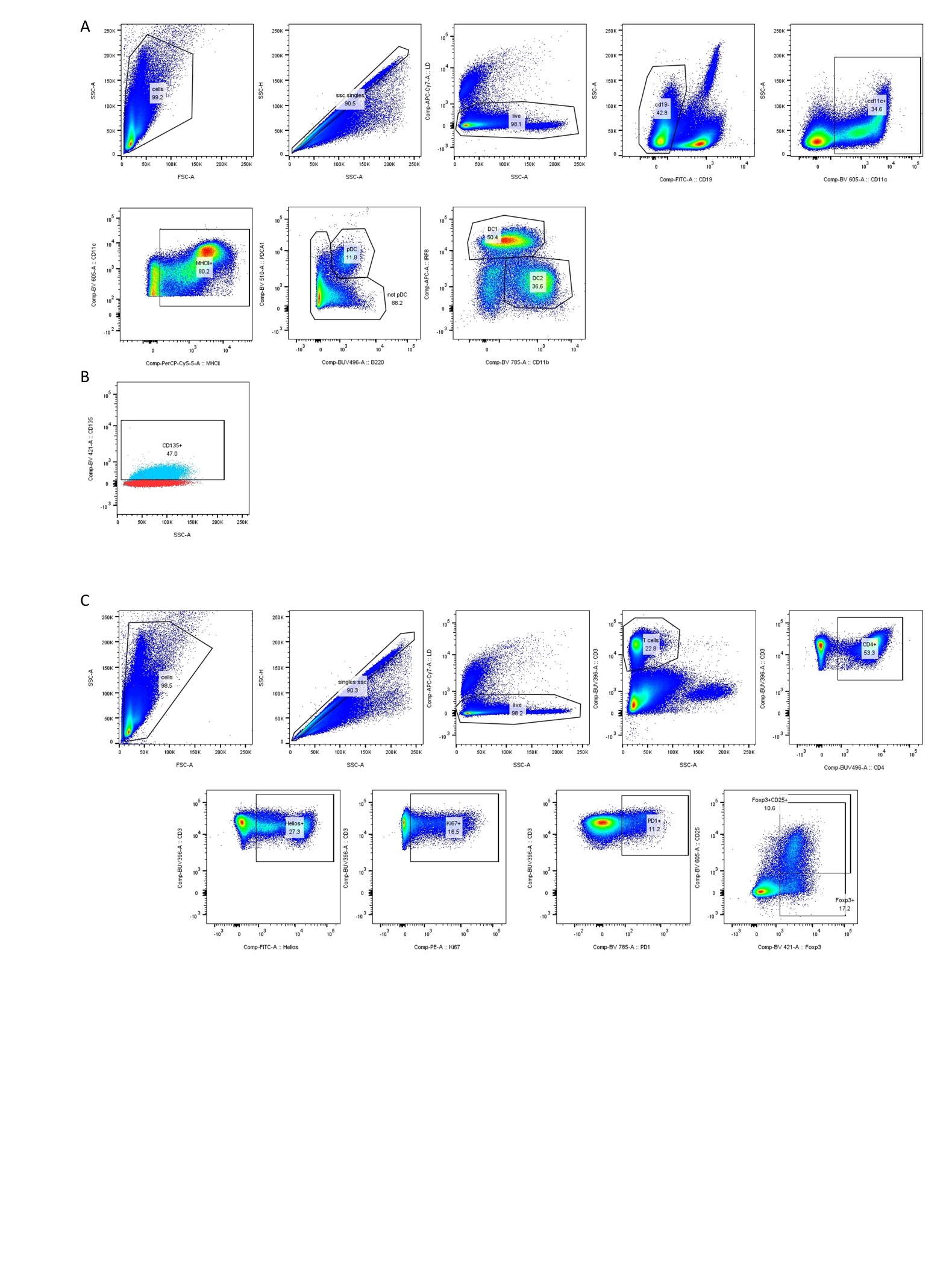

**Supplementary Figure 1: Representative flow gating for Figure 3. (a)** DC flow panel and gating. (b) Representative CD135 staining on DCs with the stained population represented in blue and the CD135 FMO in red. (c) T cell and T reg flow panel and gating.

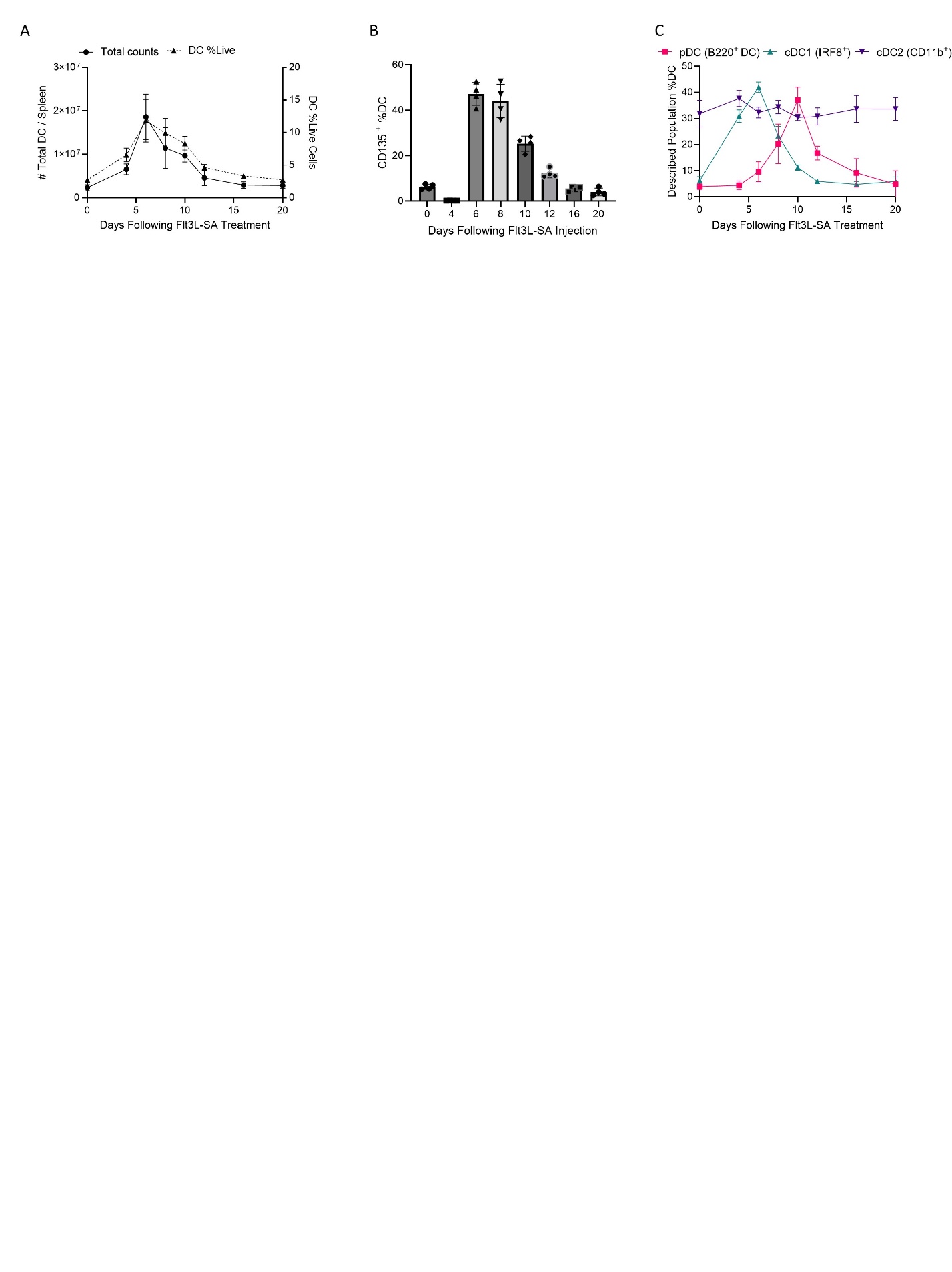

**Supplementary Figure 2: Flt3L-SA treatment expands total DCs via engagement of CD135 and skews DCs to cDC1 and pDC phenotypes. (a)** Representation of total DCs (CD11c^+^MHCII^+^) over time following Flt3L-SA treatment and (b) CD135 expression on said DCs. (c) Proportion of total DCs represented by pDC (pink), cDC1 (green), or cDC2 (purple) over time following Flt3L-SA dosing. For (a) and (c), each data point represents the mean with error bars representing SD. For (b), each data point represents one mouse with error bars for SD.

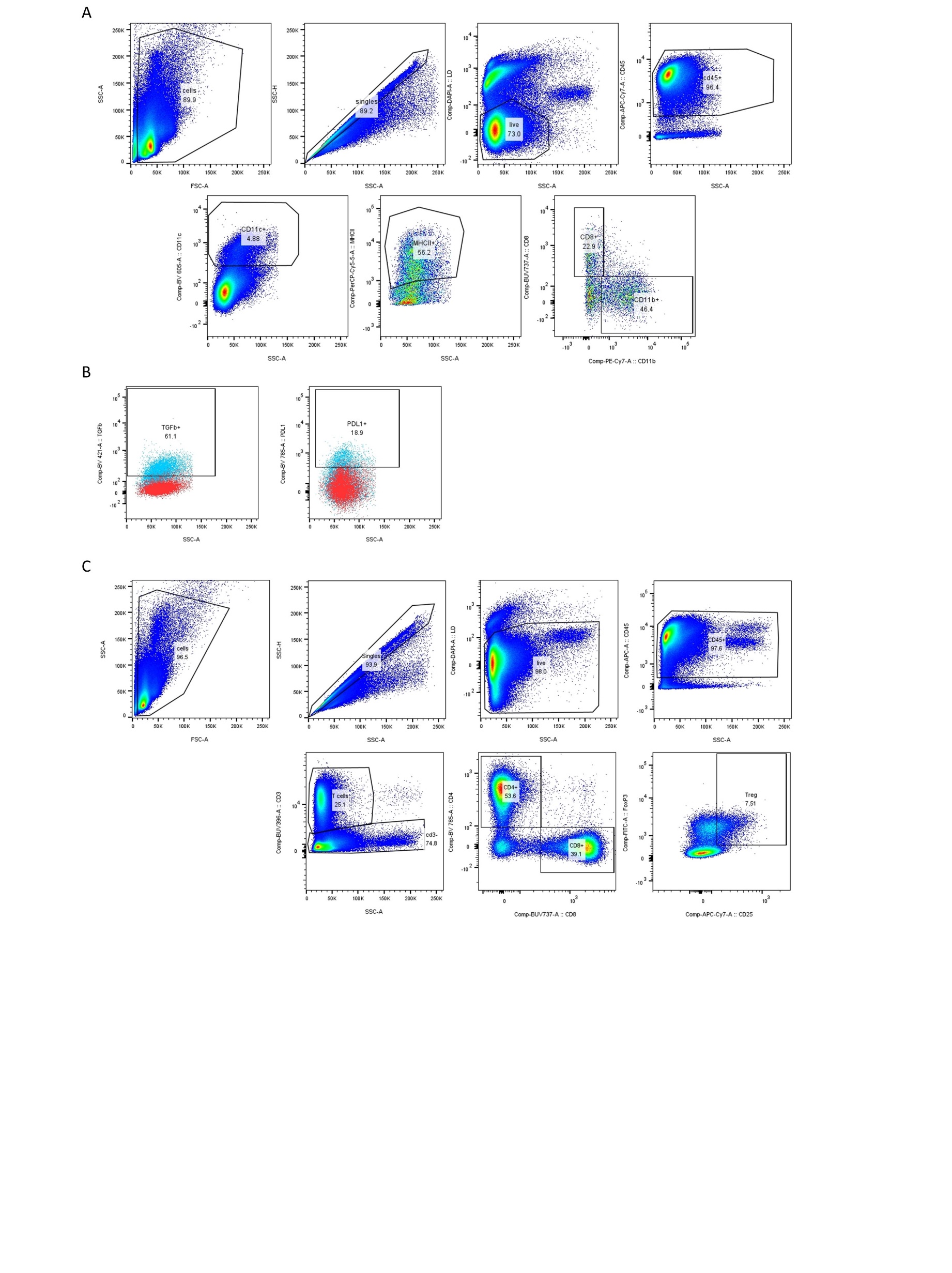

**Supplementary Figure 3: Representative gating for Figure 4. (a)** DC flow panel and gating. (b) Representative expression of LAP (left) and PDL1 (right) on total DCs with full stain in blue and FMO populations in red. (c) T cell and T reg flow panel and gating.

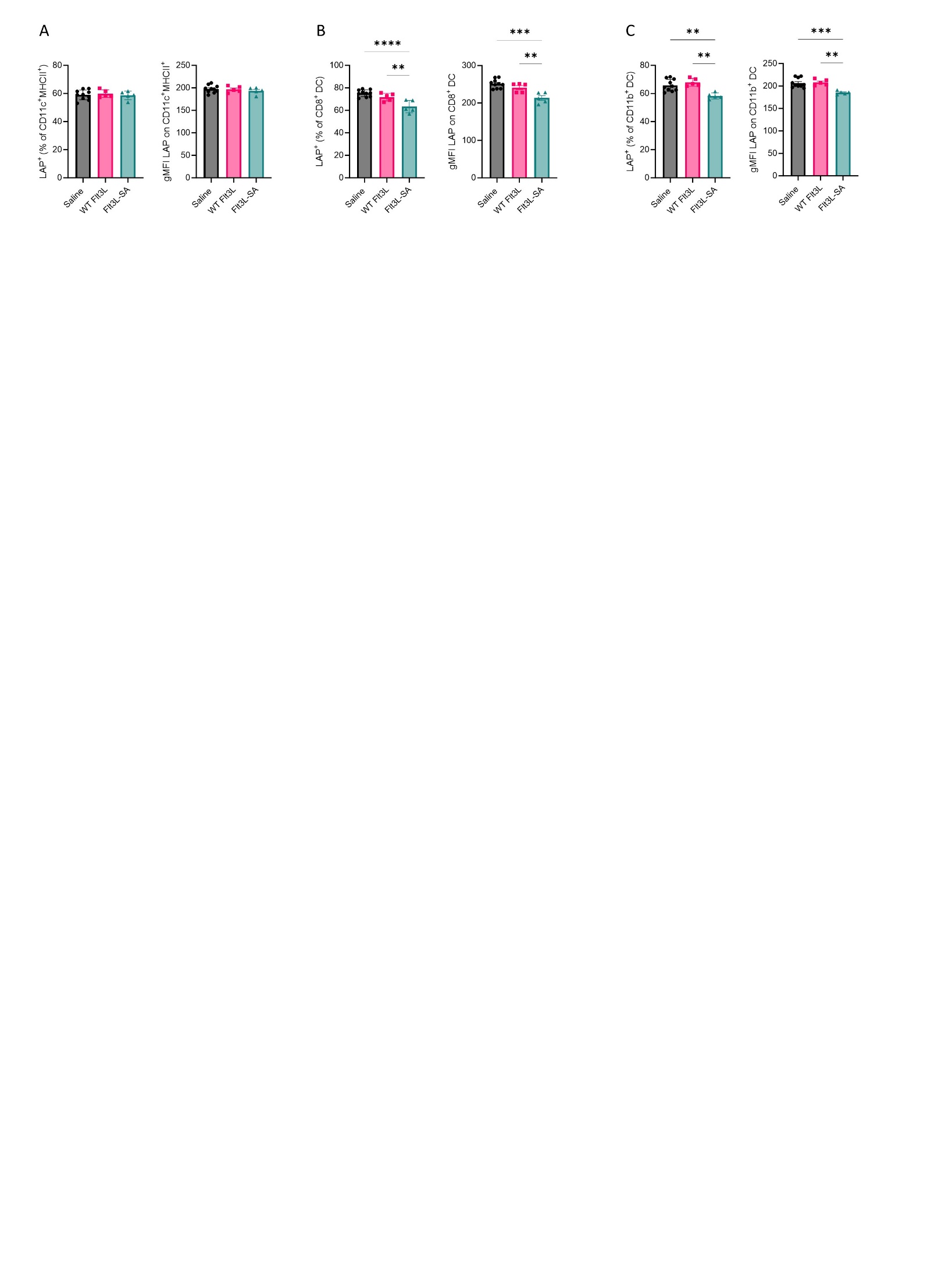

**Supplementary Figure 4: LAP expression on cDC subsets after Flt3L variant treatment. (a)** LAP expression as percent of total DCs (left) and gMFI (right) and the same measurements on (b) CD8^+^ cDC1 and (c) CD11b^+^ cDC2 cells. Each data point represents one mouse with error bars for SD. Statistics calculated via one-way ANOVA between all groups with Tukey’s multiple comparison correction. ** for p<0.01, *** for p<0.001, **** for p<0.0001.

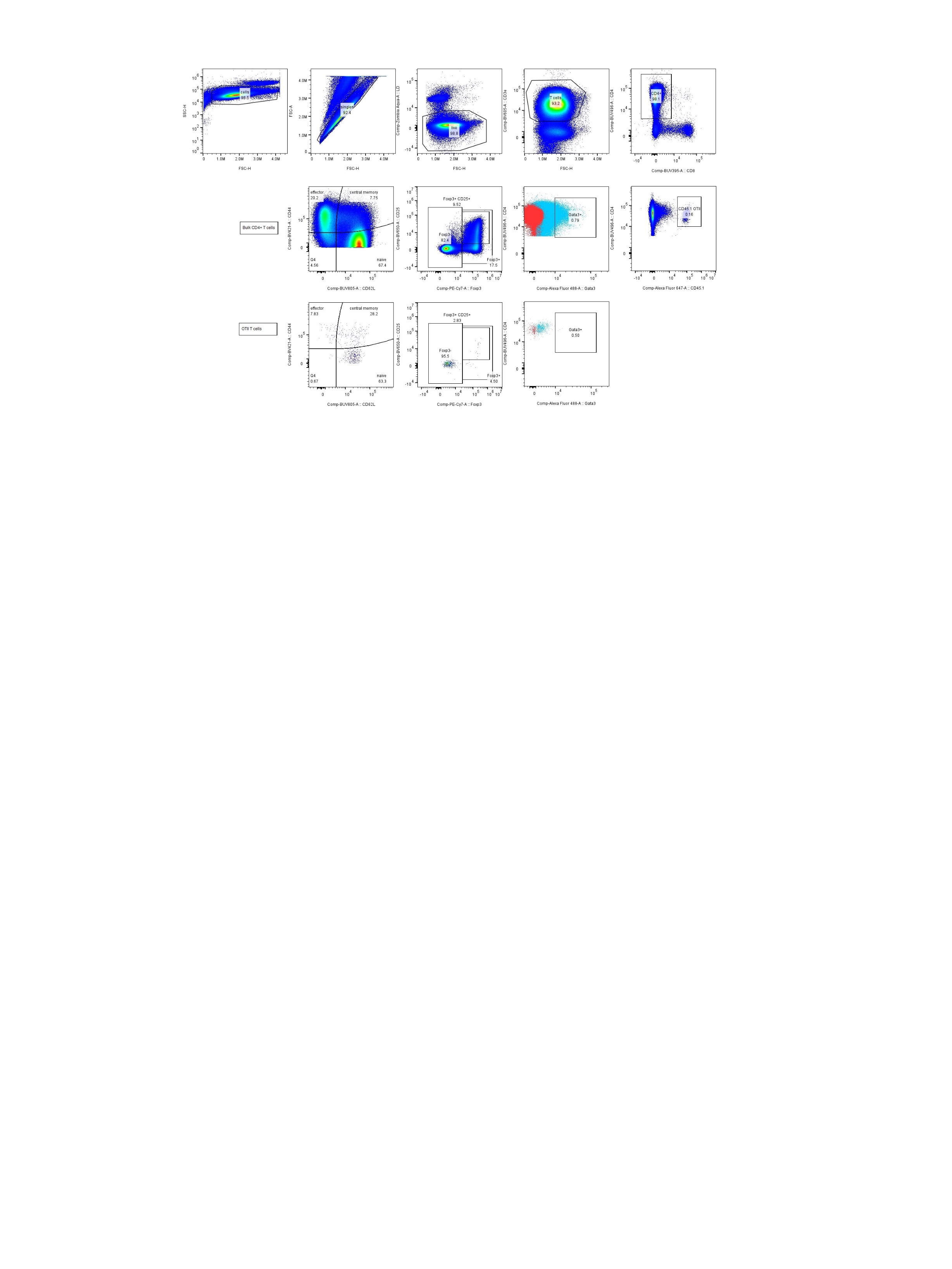

**Supplementary Figure 5: Representative gating for Figure 5.** T cell gating with full staining as representative of the gating strategy. On the Gata3 gated graphs, full stain is represented with blue dots and the Gata3 FMO is in red.

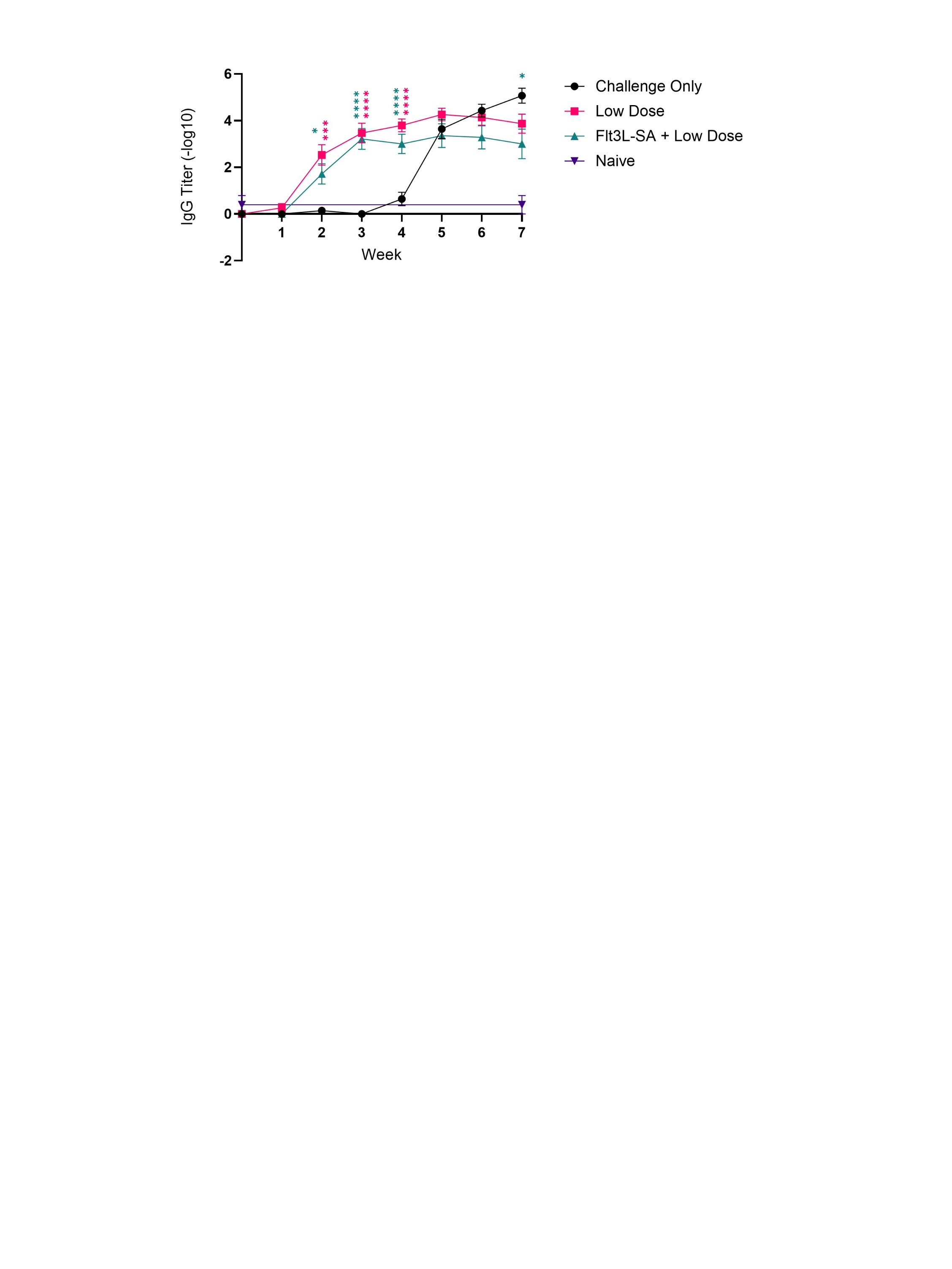

**Supplementary Figure 6: Total IgG titer against Rasburicase over entire course of study.** Each point represents the average with error bars for SEM. Significance is calculated by one-way ANOVA with Tukey’s multiple comparison at each timepoint with color denoting significance of that group compared to challenge only group. * for p<0.05, *** for p<0.001, **** for p<0.0001.

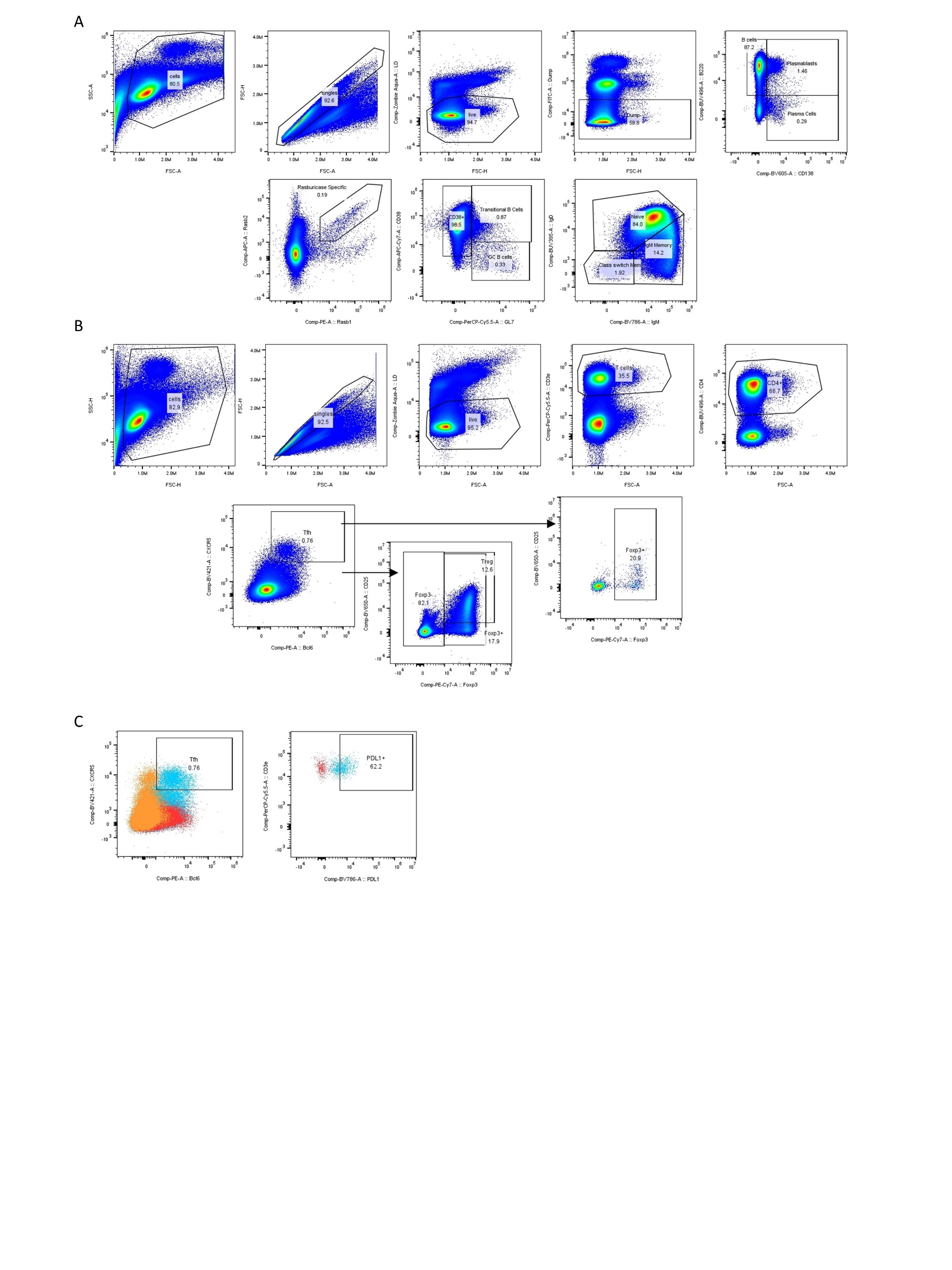

**Supplementary Figure 7: Representative gating for Figure 7. (a)** B cell representative gating on bulk populations. (b) T cell representative gating with arrows denoting the Tfh population or the “not” Tfh population moving forward for gating. (c) Tfh (left) gating on total CD4^+^ T cells with full stain represented in blue, CXCR5 FMO represented in red, and Bcl6 FMO represented in orange. PD-L1 expression (right) within the Tfr population with full stain in blue and PD-L1 FMO in red.

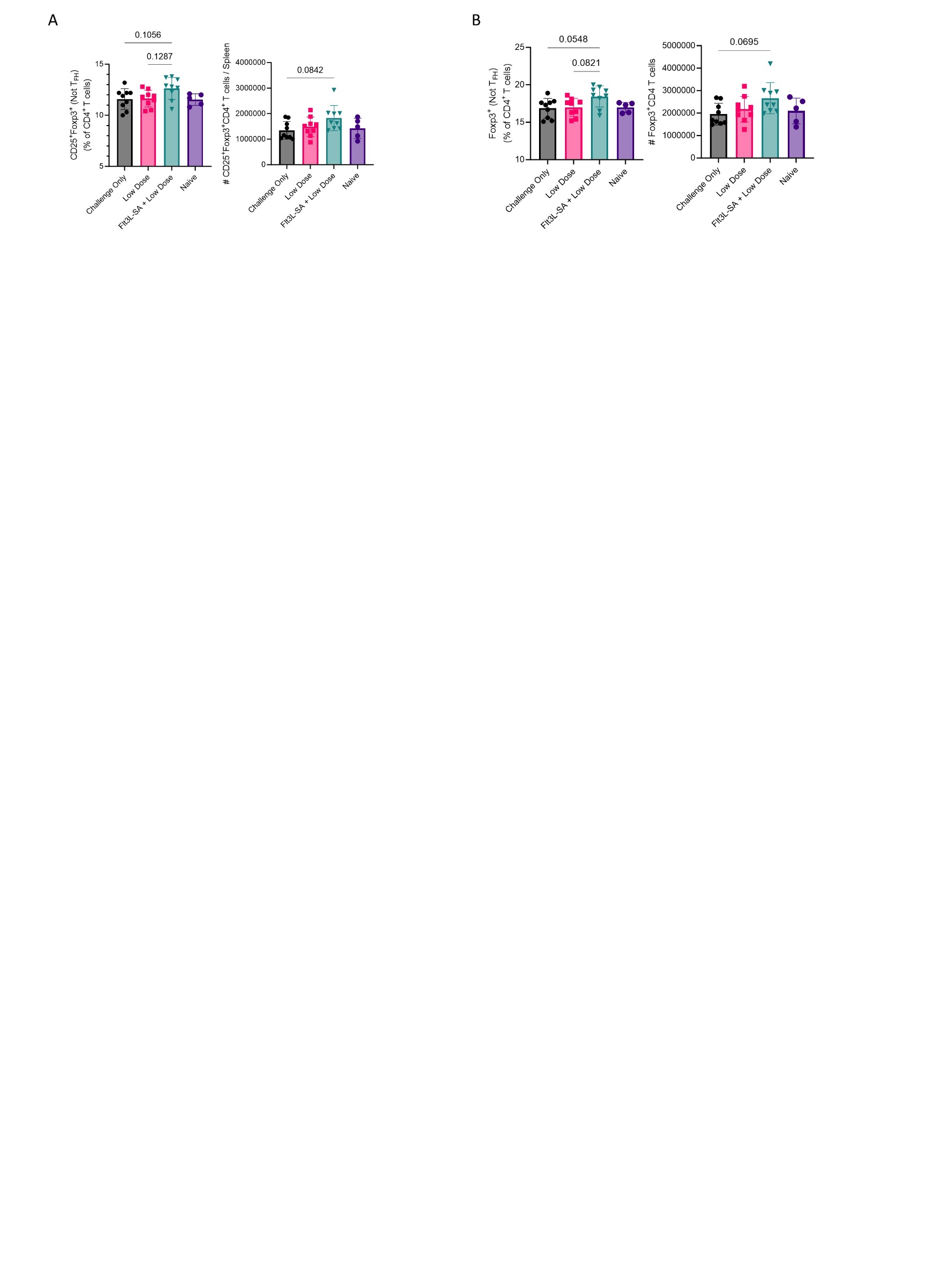

**Supplementary Figure 8: Flt3L-SA induces a trending increase in Tregs and Foxp3 expressing non-Tfh cells. (a)** Quantification of bulk Tregs (CD25^+^Foxp3^+^) and (b) Foxp3^+^ T cells in the spleen following four weeks of i.v. rasburicase challenge. Each data point represents one mouse with error bars for SD. Statistics calculated via one-way ANOVA between all groups with Tukey’s multiple comparison correction.
